# High-resolution metagenome assembly for modern long reads with myloasm

**DOI:** 10.1101/2025.09.05.674543

**Authors:** Jim Shaw, Maximillian G. Marin, Heng Li

## Abstract

Long-read metagenome assembly promises complete genomic recovery from microbiomes. However, the complexity of metagenomes poses challenges. We present myloasm, a metagenome assembler for PacBio HiFi and Oxford Nanopore Technologies (ONT) R10.4 long reads. Myloasm uses polymorphic k-mers to construct a high-resolution string graph and then leverages differential abundance for graph simplification. On real-world ONT metagenomes, myloasm assembled three times more complete circular contigs than the next-best assembler. Myloasm can make ONT and HiFi comparable for assembly: for a jointly sequenced gut metagenome, myloasm with ONT assembled more complete circular genomes than any assembler with HiFi. Myloasm recovers previously inaccessible within-species diversity; we recovered six complete *Prevotella copri* single-contig genomes from a gut metagenome and eight complete TM7 (Saccharibacteria) contigs with >93% similarity from an oral metagenome. With this improved resolution, we resolved two 98% similar *ermF* antibiotic resistance genes spreading through distinct strain-specific mobile genetic elements in a human gut.

## 1 Introduction

Metagenomics studies the collective genomic content within a community of organisms through untargeted DNA sequencing [1]. After sequencing, computational *de novo* assembly algorithms are used to reconstruct genomes. The recovery of genomes from metagenomic sequencing has led to a drastic increase in our understanding of the microbial world [2,3]. Metagenomics has enabled fundamental insights into evolution and ecology [4,5,6,7] and has emerged as a crucial tool for linking microbial communities to human health [8,9,10,11,12] or environmental health [13,14,15,16]. Many microbes may not be culturable [17], so a metagenomics approach is necessary for the discovery of new genomes. In the past decade, the emergence of long-read sequencing technologies has led to drastic improvements in isolate and metagenome recovery [18,19,20,21,22]. Nevertheless, the scale, diversity, and complexity of metagenomes remains a challenge for complete recovery of metagenomes [23].

Microbial communities can contain many organisms with distinct but similar genomes. This can be due to ongoing recombination [24,25] within a population or distinct populations of the same species [26,27]. Furthermore, horizontal gene transfer can lead to shared chromosomal DNA between species [28]. This makes metagenome assembly challenging due to the presence of highly-similar and thus repetitive sequences within a metagenome. Short-read sequencing has fundamental and practical limitations for resolving these intergenomic repeats [29,30,31], but ONT and PacBio HiFi long reads represent promising solutions. HiFi assemblers [21,20] require highly accurate reads, but this often enables more contiguous and higher-resolution assemblies compared to ONT assemblers [19]. The rapid improvement of ONT sequencing accuracies to *>* 99% [32], driven by its latest R10.4 chemistry, has the potential to close this gap [33]. However, these ONT R10.4 reads are still less accurate than HiFi, and new algorithmic techniques are required to harness their potential [34].

Modern long-read assemblers can be broadly stratified into two paradigms: string graphs [35] and de Bruijn graphs [36]. In the string graph, nodes are reads and edges are overlaps between reads. In the (node-centric) de Bruijn graph, nodes are k-mers and edges are error-free overlaps of length *k* − 1. String graphs are more powerful at resolving genomic repeats because overlaps are not restricted to length *k* − 1. However, string graphs are less computationally efficient because they require all overlaps to be explicitly computed, whereas de Bruijn graphs do not. Furthermore, errors pose a problem for exact k-mer overlaps—even a modest k-mer overlap length of 500 bp is likely to have an error for the newest ONT reads. New minimizer de Bruijn graph approaches tackle these limitations by using longer sequence contexts and error correction [37,20,38]. Nevertheless, these graphs are still losing information longer than k-mers in comparison to string graphs.

### 1.1 Our contribution

We present myloasm (**m**etagenomic nois**y lo**ng-read **as**se**m**bler), a new metagenome assembler for PacBio HiFi and ONT R10.4 long reads. In myloasm, we use a string graph approach. Unlike existing HiFi string graph approaches [39,40], we eschew error correction, which may fail for low-coverage or high-diversity populations due to the lack of reads coming from identitical genomes. We instead use polymorphic k-mers within the sample to resolve similar sequences in metagenomes (e.g., co-existing strains or conserved genomic regions). In addition, we use a graph cleaning algorithm inspired by annealing approaches from statistical physics [41] to integrate coverage and overlap information. We show that myloasm can resolve simple populations of closely related strains better than de Bruijn graph approaches, yet handle complex metagenomes better than previous string graph algorithms. This leads to improved recovery of metagenome-assembled genomes (MAGs) over previous approaches.

## 2 Results

### 2.1 Resolving overlap assembly graphs with polymorphic k-mers and a random path model

Myloasm first performs a form of reference-free variant calling via *SNPmers* (**Fig. 1A**). SNPmers are pairs of k-mers that differ by a substitution in the middle base; SNPmers have been used and defined previously in other contexts [42,43]. SNPmers capture single nucleotide polymorphisms (SNPs) through a k-mer context instead of a reference genome. Myloasm calls SNPmers by finding suitable pairs of k-mers and excluding erroneous, low-frequency k-mers. Myloasm then uses a combination of SNPmers and open syncmers [44] to index the reads.

**Fig. 1:**
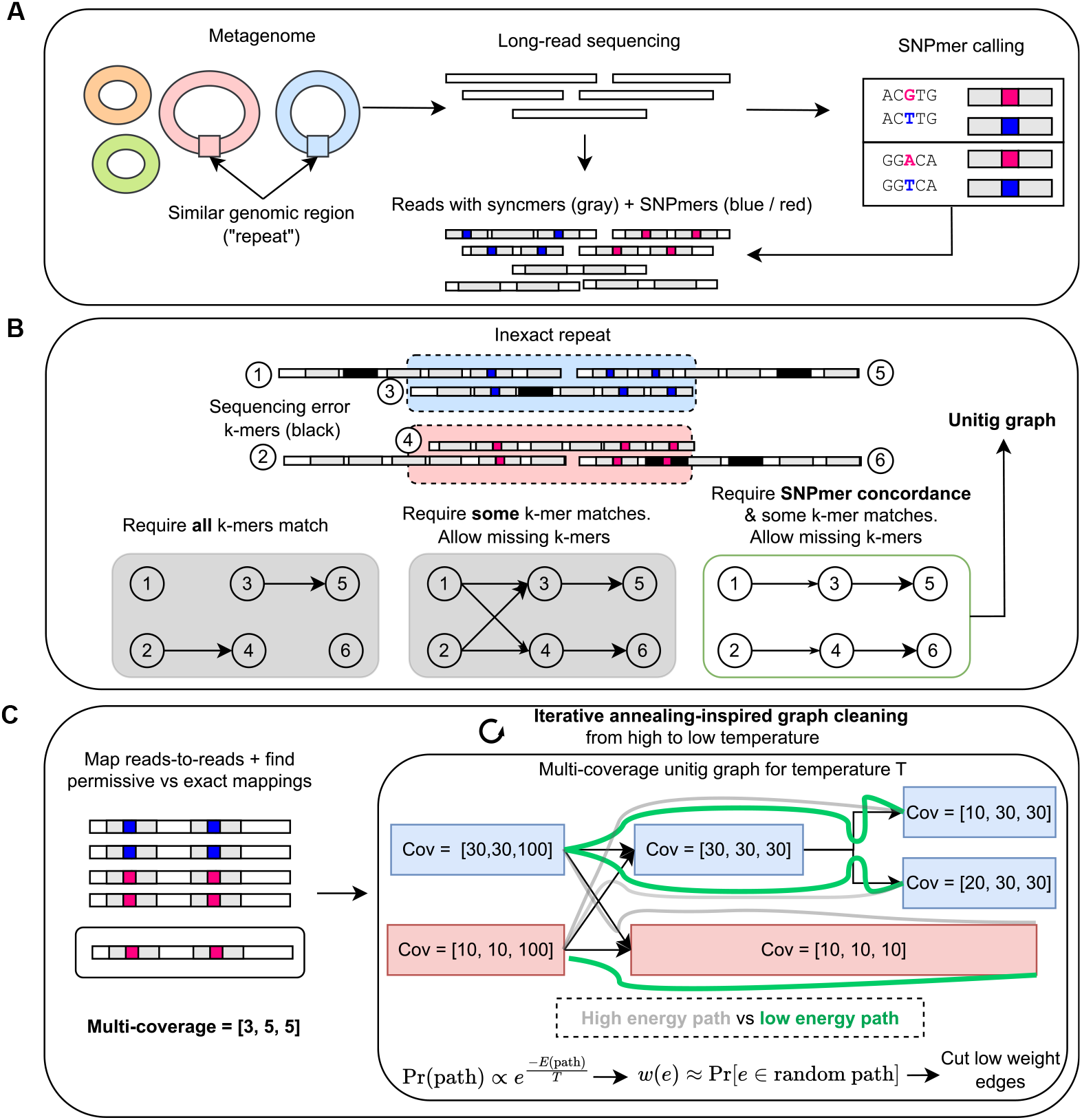
Algorithm overview of myloasm. **A**. SNPmers—k-mers that have polymorphic middle bases— are called from all sequences. Long reads are then indexed with SNPmers and open syncmers. **B**. Inexact repeats between or within genomes create multiple paths in an assembly graph. Myloasm allows for errors but penalizes for mismatched SNPmers, creating a simpler assembly graph without error correction. **C**. Reads are mapped to and coverage is calculated. Myloasm calculates several coverages at different sequence identity thresholds. These coverages are used to define a probabilistic model on paths, where paths with consistent coverage have higher probability. Edges are weighted by the path probabilities, then an iterative cleaning approach is used to clean up the graph by cutting lowly weighted edges while respecting the graph topology.

In the overlapping stage (**Fig. 1B**), myloasm finds exact open syncmer matches and then performs chaining [45,46] to find overlaps. Then, myloasm matches SNPmers while *ignoring the middle base*. That is, myloasm allows matching of polymorphic SNPmers. Myloasm then performs chaining again, but this time on the SNPmers. This double chaining procedure can estimate a true sequence divergence (and thus, sequence identity) for the mapping through a statistical model of mutation and sequence error. Myloasm uses this estimate of sequence identity to build a string graph. This step is permissive for missing syncmer or SNPmer hits between reads; error correction is not required. However, it is strict for *mismatched SNPs* (i.e., mismatched middle bases) across SNPmers (**Fig. 1B**, bottom). After transitive reduction, non-branching paths are collapsed into unitigs.

Myloasm uses differential abundance within a metagenome to resolve the unitig graph (**Fig. 1C**). Myloasm first performs read-to-read mapping and calculates depth of coverage. Importantly, myloasm calculate coverages at different identity cutoffs for mapping. We found this to be necessary under extensive strain variation, where few perfect mappings may exist. Myloasm then uses the coverage to simplify the graph. We describe this below.

Myloasm weighs edges by their likelihood to appear under a random path model. Paths with coverage discordance and containing small overlaps with high divergence will be assigned a lower probability. We parameterize this model by a temperature parameter that controls the strictness of the penalty. Myloasm then cuts edges with small expected value of appearing in a random path. Finally, myloasm removes tips and bubbles [47], and collapses paths into unitigs. However, we do not use a fixed temperature value. Instead, we iterate this cleaning procedure from high to low temperature. The intuition is that the initial stages (high temperature) cut only confident edges, leading to longer unitigs. These longer unitigs have more robustly estimated edge weights. Then, myloasm can prune edges more aggressively in later steps (lower temperatures). After cleaning, the final unitigs are polished and output to the user as contigs.

### 2.2 High-resolution assemblies on a concatenated mock ONT R10.4 community

We constructed a mock long-read metagenome with 48 genomes sequenced to 19 Gbp and basecalled with both sup and hac ONT R10.4 reads. This was done by concatenating two real mock metagenomes and 14 isolate sequencing runs [32]. We focus on the results for the sup basecalled reads **Fig. 2**; hac results are shown in **Supplementary Fig. 1**.

**Fig. 2:**
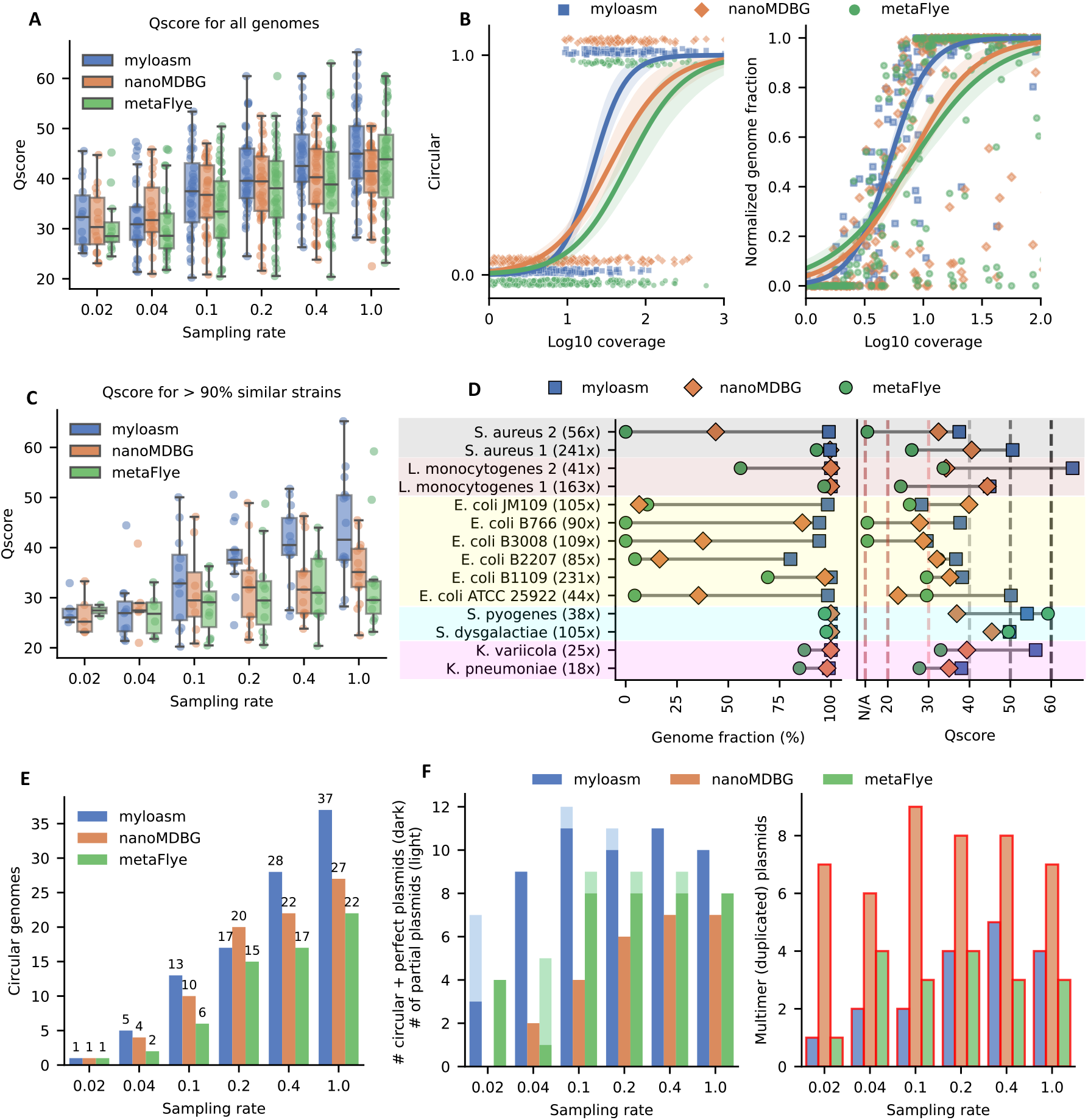
Results on a concatenated ONT R10.4 mock metagenome with 48 genomes consisting of 14 isolates and 2 mock metagenomes. **A**. Qscores (− log_10_(indel rate + substitution rate) for all 48 genomes on various downsampling rates. **B**. Circular and complete genomes and fraction of genome recovered as a function of coverage over all downsampling rates. Logistic regression was run over the dataset with confidence intervals found by bootstrapping over 1000 iterations. **C**. Qscore restricted to only genomes with a *>* 90% ANI strain within the metagenome. **D**. Qscore and genome fraction for genomes with a *>* 90% ANI strain within the metagenome on the 1.0 sampling rate dataset. Depth of coverage is shown to the right of the genome name. **E**. Number of circular complete prokaryotic genomes. **F**. Plasmid results. Left: dark bars show the number of circular + complete plasmids and light bars shown non-circular but complete plasmids. Right: number of assembled plasmids contigs that are *>* 10% longer than the reference plasmid genome, indicating possible multimeric assembly. Box plots show the median (middle line), upper and lower quartiles (lower and upper box limits) and 1.5 times the interquartile range (whiskers).

Compared to metaMDBG (hereafter assumed to use the nanoMDBG mode [38] on ONT data) and metaFlye [19], myloasm has the highest median Qscores (i.e., −10 log_10_(error rate)) on five of the six datasets after downsampling (**Fig. 2A**). At low coverage, myloasm is more likely to assemble a complete circular genome and recover more of the genome than other assemblers (**Fig. 2B**). Myloasm circularized 92% of genomes with coverage *>* 50, whereas metaFlye and metaMDBG circularized 59% and 65% respectively. We show the number of misassemblies in **Supplementary Fig. 2**. The number of misassemblies decreased as a function of coverage, but we found that no method was consistently the best or worst.

To better understand assembly behavior for closely related strains, we focused our analysis on groups of genomes with *>* 90% average nucleotide identity (ANI) to each other. This gave five groups: 90.86% ANI between *Streptococcus pyogenes* and *Streptococcus dysgalactiae*, 94.74% ANI between *Klebsiella pneumoniae* and *Klebsiella variicola*, 95.37% between two *Listeria monocytogenes* strains, 98.86% between two *Staphylococcus aureus* strains, and 96.81-99.53% ANI between six *Escherichia coli* strains.

Myloasm had a median Qscore of 41.5 compared to 35.1 of metaMBG and 28.6 of metaFlye (**Fig. 2C**) for the non-downsampled dataset on these five groups of genomes. Importantly, high Qscore does not imply high genome recovery (**Fig. 2D**). The two *Streptococcus* strains of ≈ 91% similarity were recovered by all three methods. However, for the *L. monocytogenes* and *Klebsiella* genomes at ≈ 95% identity, metaFlye recovered *<* 90% of the genome for three of the four strains. MetaFlye did recover *>* 95% for one of the *L. monocytogenes* genomes, but its Qscore was *<* 25, indicating serious polishing errors when strain heterogeneity is present. The two 98.89% similar *S. aureus* strains could only be fully recovered by myloasm. The low-coverage *S. aureus* strain posed a problem for metaMDBG (*<* 50% genome recovered) and metaFlye (no contigs of length *>* 100 kb). Thus, myloasm can recover high-quality strain-level genomes, even under varying coverages.

The six *E. coli* genomes were particularly challenging. Five were all *>* 98.3% similar, with B1109 and JM109 being 99.53% similar. Myloasm recovered *>* 75% of each *E. coli* genome. MetaMDBG only recovered *>* 75% of 2/6 genomes and metaFlye could not recover *>* 75% of any genome. Myloasm had the highest Qscore on 5/6 *E. coli* genomes; we found that a single circular contig was assembled for JM109, but it had strain switching errors, leading to low Qscore. Myloasm was the only method that recovered the low-coverage ATCC strain and had a Qscore of *>* 50 (*<* 1 error per 100kb). This strain had relatively lower coverage to the other *E. coli* s, but it was more dissimilar to the other *E. coli* s (96.81% similar). This again highlights myloasm’s ability to recover low-coverage strain-level genomes relative to the other assemblers.

Myloasm had the most circular single-contig prokaryotic genomes (**Fig. 2E**) across 5/6 sampling rates and perfectly recovered plasmids across all sampling rates (**Fig. 2F**). A large number of duplicated plasmids were present in the output assembly. This has been corroborated by other long-read assembly benchmarks [48]. We found that some reads spanned a single plasmid genome multiple times—one read spanned a *S. aureus* plasmid exactly six times, and all methods failed to correctly recover this plasmid. This multimer phenomenon has been described before for nanopore sequencing [49]. Nevertheless, myloasm and metaFlye were the most precise across all datasets, with 18 in total for each, whereas metaMDBG had 45 duplicated plasmids.

### 2.3 Enhanced metagenome-assembled genome (MAG) recovery on diverse metagenomes

We next benchmarked myloasm, metaMDBG, and metaFlye on six real R10.4 ONT metagenomes [50,51,52] and six real PacBio HiFi metagenomes [53,54,55,56,20,57] (see **Supplementary Table 1**). We also benchmarked hifiasm-meta but only for the HiFi datasets. We show MAG recovery results in **Fig. 3A** for single-sample assembly and binning with CheckM2 [58] evaluation.

**Fig. 3:**
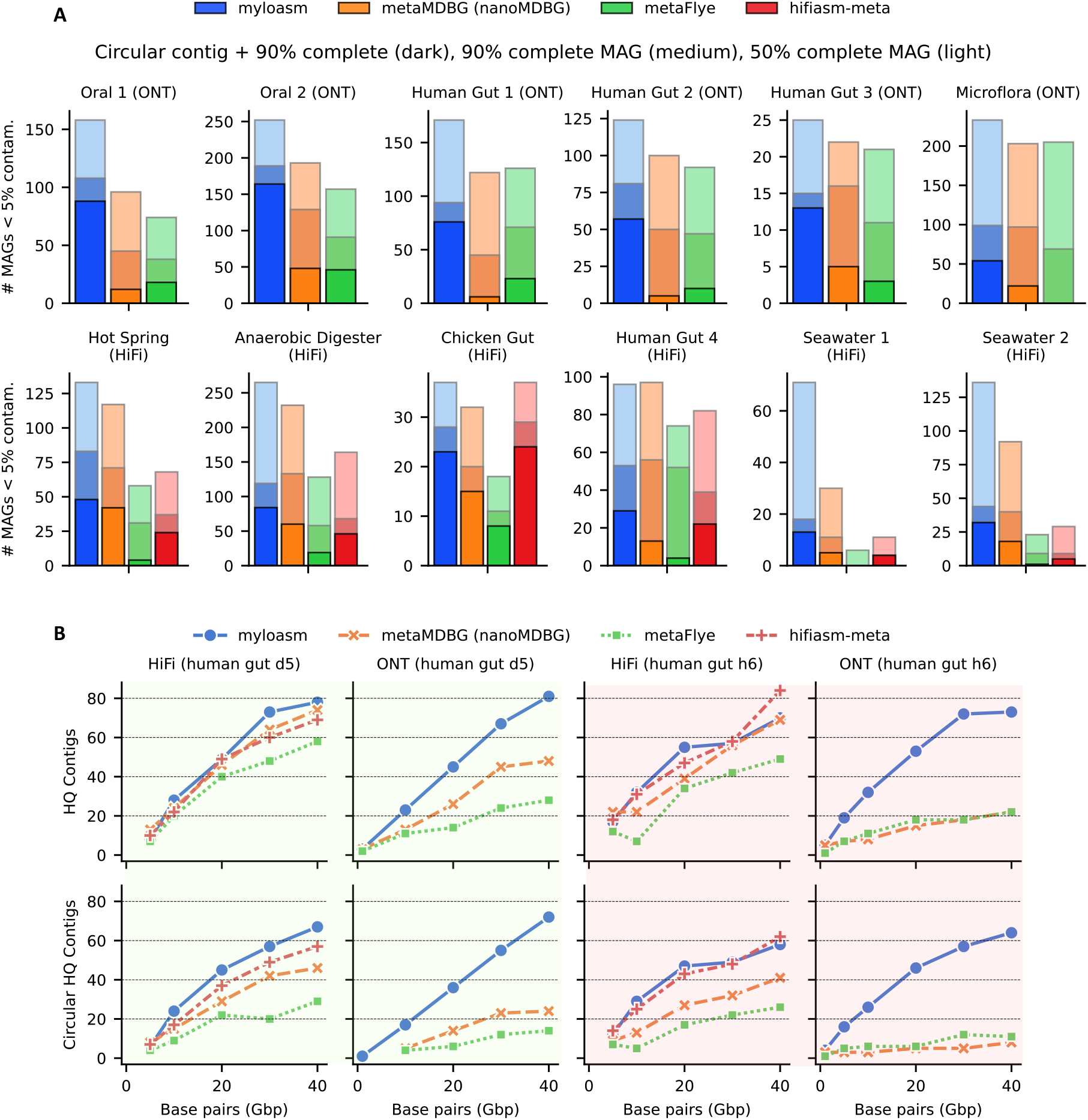
MAG and contig recovery results for real metagenomes. **A**. MAG recovery after assembly and binning. Dark bars represent circular contigs with *>* 90% completeness and *<* 5% contamination. Medium bars represent non-circular MAGs with *>* 90% completeness and *<* 5% contamination. Light bars represent MAGs with *>* 90% completeness and *<* 5% contamination. NanoMDBG is used for ONT reads whereas metaMDBG is used for HiFi reads. **B**. Top: number of high-quality contigs with *>* 90% completeness and *<* 5% contamination on two different gut samples from Minich et al. sequenced with HiFi and ONT R10.4 data. Bottom: the same datasets but counting only circular high-quality contigs. Backgrounds are colored by gut sample.

Myloasm had particularly strong performance for recovering circular, complete contigs on ONT data. Myloasm recovered *>* 3 times more circular, complete contigs for the oral datsets than the next best method. On the three gut datasets, myloasm recovered *>* 3, 5, and 2 times more circular complete contigs than the other two methods. On the complex soil ONT dataset (“Microflora”) [52] with *>* 100 Gbp, myloasm recovered 54 circular contigs versus 23 for metaMDBG and 0 for metaFlye. Myloasm also recovered more non-circular high-quality MAGs (*>* 90% complete and *<* 5% contaminated) on 5/6 ONT datasets and more *>* 50% complete and *<* 5% contaminated MAGs on 6/6 datasets. Interestingly, a single circular contig made up *>* 80% of myloasm’s high-quality MAGs on 4/6 ONT datasets; no other assembler had even 50%. This hints that, with long enough reads, a single circular contig is starting to become the expectation for high-quality MAGs.

For HiFi reads, myloasm recovered the most circular, complete contigs for 5/6 datasets but had slightly less than hifiasm-meta for the chicken gut (24 for hifiasm-meta vs 23 myloasm). Myloasm recovered the most high-quality non-circular MAGs on 4/6 datasets as well. The results diverged particularly on complex seawater metagenomes. For example, metaFlye did not recover a single high-quality MAG from the Seawater 1 dataset [55] whereas myloasm had 18. Hifiasm-meta struggled on these complex metagenomes as well. Hifiasm-meta recovered *<* 4 times less high-quality MAGs on the Seawater 2 dataset [54] compared to myloasm and metaMDBG. Overall, myloasm can recover more complete and contiguous MAGs on a variety of sequencing technologies and environments.

### 2.4 Timing and memory benchmarks on real data

Myloasm achieved the fastest assembly times on 9/12 datasets (**Supplementary Table 2**). Myloasm was faster than metaFlye on every dataset and faster than metaMDBG on 10/12 datasets. Myloasm used less peak memory than metaFlye on 10/12 datasets (0.92 *×* as much peak RAM on average) and hifiasm-meta on 4/6 datasets (0.85*×* peak RAM on average). However, myloasm consistently required more memory than metaMDBG (average 6.86*×* higher peak RAM usage), reflecting the efficiency of metaMDBG’s sparse minimizer approach. The microflora dataset was an outlier where myloasm’s RAM was *>* 2.5*×* metaFlye. We found that the in-memory k-mer counting step took ≈ 600 GB of RAM due to the high diversity of this metagenome; the rest of the algorithm took *<* 250 GB. Besides high complexity metagenomes, myloasm provides substantial runtime improvements with competitive memory usage.

### 2.5 Myloasm makes ONT sequencing competitive with HiFi for the human gut

We compared all assemblers across HiFi and ONT R10.4 using two gut datasets of Minich et al. [50]. (**Fig. 3B**) with both types of sequencing. Minich et al. found that HiFi still outperformed ONT for metaMDBG and metaFlye, producing 2.4 times more circular complete genomes than ONT. With myloasm, this is no longer the case. With ONT, myloasm generated an average of 104.3% (h6) and 88.7% (d5) of the best HiFi assembler’s circular contigs across all subsamples. For the same metrics, metaMDBG achieved only 13.1% (h6) and 32% (d5); metaFlye achieved only 22.3% (h6) and 18% (d5).

Assembly critically depends on the read length distributions, which differs between ONT and HiFi for the same biological samples. For sample d5, the N50 and N10 read lengths were N50 = 11 kbp, N10 = 19 kbp (HiFi) and N50 = 10 kbp, N10 = 26 kbp (ONT). Thus, ONT has more very long reads compared to HiFi. For h6, N50 = 12 kbp, N10 = 17kbp (HiFi) and N50 = 6 kbp, N10 = 33 kbp (ONT). The ONT h6 sample had a heterogenous length distribution that was approximately 2*×* smaller for N50 than HiFi, yet myloasm recovered more circular contigs on average with ONT than HiFi. Thus, myloasm can leverage the presence of very long reads, closing the gap between HiFi and ONT for high-quality contig recovery for human gut metagenomes.

### 2.6 Contig and MAG-level contamination assessment

For the same 12 datasets as previously described, we ran CheckM2 on all contigs of length *>* 500 kbp and show the results in **Fig. 4A**. We show the analogous results for MAGs in **Supplementary Fig. 3**. On the six ONT datasets, myloasm produced more *>* 50, 75, and 90% complete contigs with low contamination (*<* 5%) than every other assembler. For HQ contigs (*>* 90% complete and *<* 5% contaminated), myloasm output 3.9*×* more *>* 90% HQ contigs than metaFlye and 2.8*×* more than metaMDBG on average. For HiFi, metaMDBG was more competitive; myloasm produced 1.3*×* more HQ contigs, which was a relatively smaller increase than the ONT case. Thus, myloasm consistently captures more complete genomes as a single contig than the other assemblers.

**Fig. 4:**
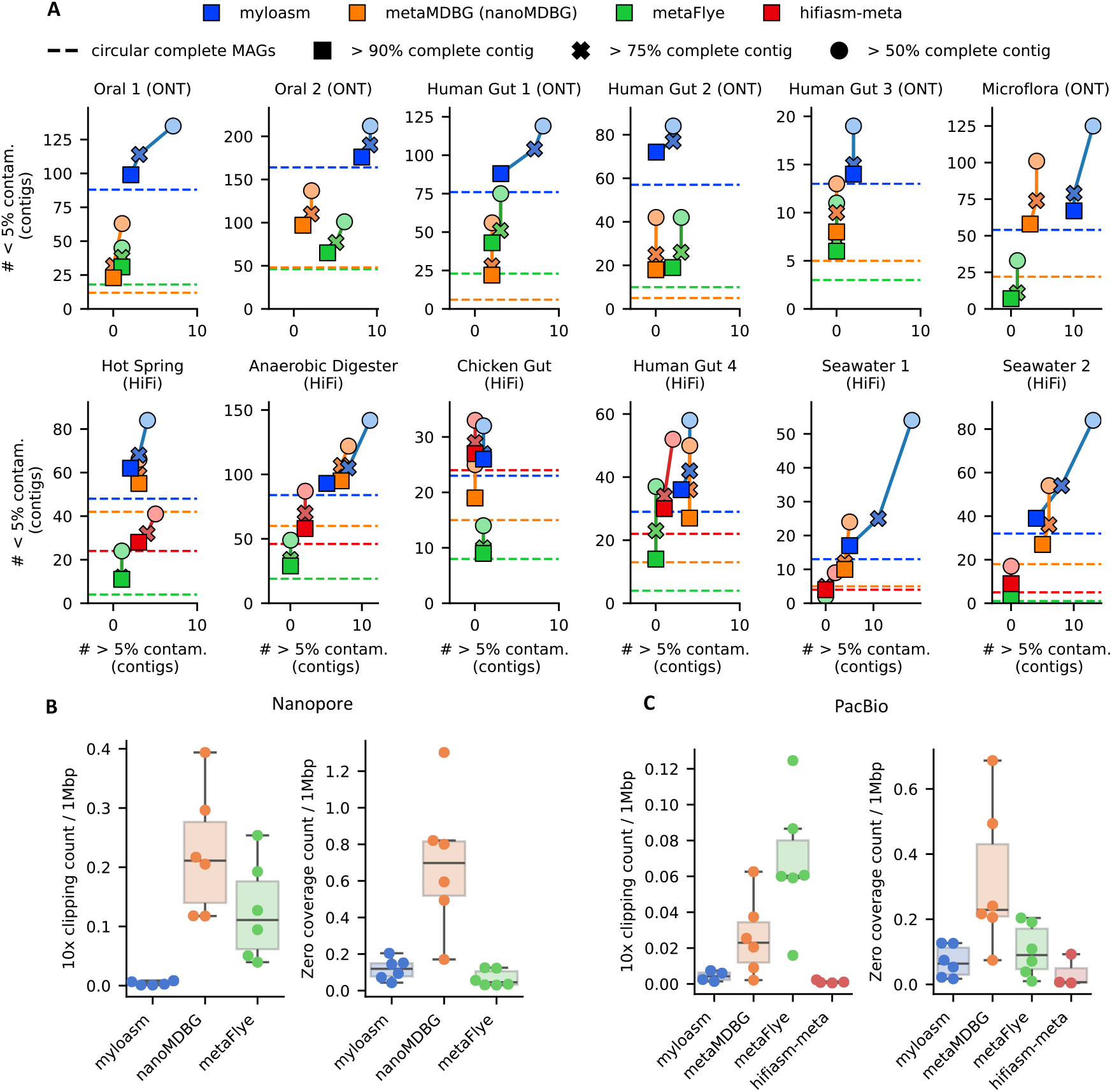
Quality control assessment for contigs. **A**. CheckM2 evaluation on contigs of length *>* 500 kbp from the same datasets as **Fig. 3**. The dashed line indicates circular complete contigs with *>* 90% contamination and *<* 5% completeness. **B**. The number of regions per 1 Mbp with *>* 10*×* coverage and every alignment at the base was soft clipped. **C**. The number of *>* 500 bp regions with zero coverage after aligning to the contigs.

As expected with this greater sensitivity, myloasm also produces more contigs that had *>* 5% contamination (denoted as high contamination). Of the *>* 90% complete contigs across all datasets, myloasm produced 789 with low contamination (*<* 5% contamination) and 45 with high contamination, with a low/high ratio of 17.5. MetaFlye had a low/high ratio of 11/236 = 21.4 and metaMDBG had 29/459 = 15.8. Thus, myloasm had competitive relative contamination for high-completeness contigs.

For *>* 50% complete contigs, myloasm obtained 1.5*×* more contigs than metaMDBG and 2.6*×* more than metaFlye. However, the low/high contamination ratio was worse for myloasm (12.5) than metaMDBG (21.5) and metaFlye (27.5). However, for *>* 50% complete *bins*, not contigs, we found that myloasm actually had a better low/high contamination ratio: the ratios were 6.1 (myloasm), 5.8 (metaMDBG), 5.8 (metaFlye). Furthermore, myloasm recovered 1.27*×* and 1.73*×* more *>* 50% complete and low-contamination bins than metaMDBG and metaFlye across all datasets.

### 2.7 Reference-free assessment of assembly quality

We examined read mapping patterns against contigs to assess quality using the same analysis as in a recent study [59] (**Fig. 4B, C**). We measured the number of regions in the assembly with (1) positions with only soft-clipped alignments yet *>* 10*×* depth of coverage (read-clipped regions; RCRs) and (2) no read support in a *>* 500 bp window (zero-coverage regions; ZCRs). These regions indicate likely local errors in the assembly. Long-read assemblers can erroneously call non-circular contigs as circular. We attempted to quantify the number of erroneously or prematurely circularized contigs but ran into several issues (see Section 4.19). Therefore, we defer the assessment of poor circularization to next section on plasmids and viruses.

We found fewer read-clipped regions (RCRs) per 1 Mbp in myloasm on average (0.005 for ONT; 0.004 for HiFi) compared to metaFlye (0.12 for ONT; 0.06 for HiFi) and metaMDBG (0.22 for ONT; 0.026 for HiFi). Howver, hifiasm-meta was the most accurate for HiFi data (0.001 RCRs per 1 Mbp). For zero-coverage regions (ZCRs) per 1 Mbp, myloasm + ONT was slightly worse (0.11) than metaFlye + ONT (0.06), but myloasm was better for HiFi (0.07 ZCRs for myloasm vs 0.10 ZCRs for metaFlye). Hifiasm-meta was also the best (0.03 ZCRs) on HiFi, and metaMDBG was the worse on both technologies (0.70 ZCRs and 0.32 ZCRs for ONT and HiFi respectively).

### 2.8 Recovery of plasmids and viruses

We evaluated the recovery of plasmids and viruses (**Fig. 5**) by predicting plasmid/viral contigs with geNomad [60] and then marking a contig as duplicated if the k-mer multiplicity is *>* 1.25. We focus on duplications as a measure of structural assembly error because (1) duplications are common for long-read plasmid misassemblies [48] and (2) they can be estimated without a reference genome. Plasmid/viral prediction software can have false positives, so we restricted analysis to contigs with a plasmid conjugation gene (including mobilization genes) detected by genomad or circular plasmids (**Fig. 5A**). For viruses, we used CheckV on predicted contigs to measure the sensitivity of high-quality (HQ) virus recovery (**Fig. 5B)**.

**Fig. 5:**
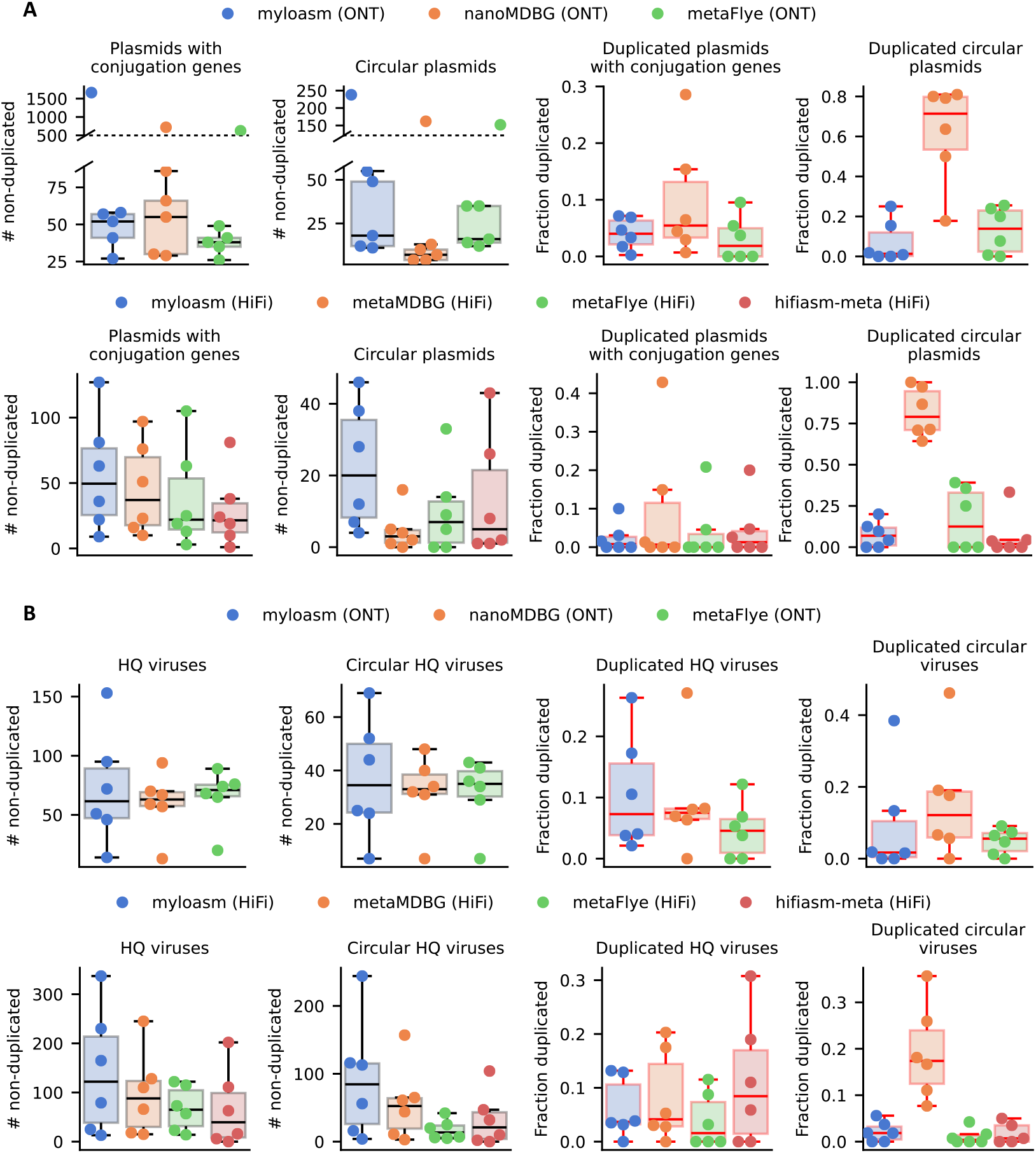
Plasmid and virus recovery results for the six ONT and six HiFi datasets. **A**. Plasmid results for the ONT datasets (top) and HiFi datasets (bottom). The two left plots only count putative plasmid contigs that pass a k-mer filter for repetitiveness. The right two plots show the fraction of plasmids with conjugation or mobilization genes or circular plasmids that were deemed to be overly repetitive and hence duplicated. **B**. Viral results for ONT (top) and HiFi (bottom). The two left plots count non-duplicated high-quality (HQ) contigs as assessed by the same k-mer filter and CheckV. The right two plots show the fraction of HQ viruses and circular viruses that were deemed duplicated using the k-mer filter.

For ONT and plasmids, myloasm recovered comparable non-duplicated plasmids to metaMDBG (median 54.5 for myloasm, 60.5 for metaMDBG, and 39.5 for metaFlye). The complex microflora dataset was an outlier with 1664 contigs with conjugation genes recovered by myloasm; 85% of these contigs had depth of coverage ≤ 2 as estimated by myloasm’s internal coverage calculator, thus indicating some degree of fragmentation. For HiFi, myloasm recovered more non-duplicated plasmids (median 49.5 for myloasm) than the other three methods (22 for metaFlye, 37 for metaMDBG, and 21.5 for hifiasm-meta). For circular and non-duplicated plasmids, myloasm had the best recovery (median 23) compared to metaFlye (14) and metaMDBG (4.5). MetaMDBG performed poorly for circular plasmids: a median of 71.4% of circular plasmids were duplicated for ONT and 79.0% for HiFi. Myloasm had the lowest duplication rate (median of 1.3% vs 13.8% for metaFlye) for ONT and second-best for HiFi (median of 6.9% vs 1.9% for hifiasm-meta and 12.5% for metaFlye).

All three methods recovered similar numbers of HQ viruses for ONT; each method was within *±*15.5% for median viruses recovered. However, for HiFi, myloasm recovered more HQ viruses (median of 122 non-circular; 84.5 circular) than metaMDBG (88 non-circular; 52.5 circular), flye (65 non-circular; 13.5 circular) and hifiasm-meta (39.5 non-circular; 21 circular). The discrepancy was notable on the Seawater 2 dataset, where myloasm recovered 244 circular HQ and non-duplicated viruses, whereas metaFlye recovered 42 (*>* 5.8 fold difference). MetaMDBG’s rate of circular duplication was still higher for both ONT (12% median vs 5.6% for metaFlye and 1.7% for myloasm) and HiFi (17.4% vs 0.4% for metaFlye, 0.4% for hifiasm, and 1.7% for myloasm). Overall, myloasm improves recovery of putative circular plasmid and viral contigs across a range of metagenomes and technologies.

### 2.9 Myloasm reveals within-species heterogeneity and strain-specific dynamics in ONT metagenomes

We investigated myloasm’s ability to recover similar coexisting genomes within real communities. For myloasm’s assemblies on the six ONT datasets, we counted the high-quality (HQ) contigs (i.e., *>* 90% complete and *<* 5% contaminated) with *>* 70% aligned fraction (AF) and *>* 93% ANI to another contig within the assembly as a rough species-level similarity threshold. For the three gut metagenomes, 26 out of 174 high-quality contigs passed the threshold (14.9%). The two oral metagenomes had higher heterogeneity, with 157 out of 275 (57.1%) passing the threshold. For the microflora soil dataset, 6 out of 65 (9.2%) passed the threshold.

We first highlight a gut metagenome (Gut 2) with interesting patterns of within-species diversity (**Fig. 6A-C**). Phylogenetic analysis of single-contig, high-quality (HQ) genomes (**Fig. 6A**) showed more comprehensive recovery with myloasm (72 genomes and 10 phyla) than nanoMDBG (18 genomes and 8 phyla) and metaFlye (18 genomes and 8 phyla). Myloasm assembled six *Prevotella copri* [61,62] (also known as *Segatella copri*) single-contig HQ genomes in this sample (**Fig. 6A, B**), with four of them circular. All other species in the Bacteroidota phylum only had one genome recovered. The *P. copri* genomes had length 4.0-4.3 Mbp with mean completeness 99.6% and contamination 1.2%. All genomes had 73-78% AF and 97.76-98.19% ANI to each other. Interestingly, we also assembled five circular elements between 127 and 155 kbp that had sequence homology to known unusual extrachromosal elements of *P. copri* [61]. Four out of six of these circular elements had mean depth of coverage within one of a *P. copri* genome and large shared regions, allowing us to infer host linkage (**Supplementary Fig. 4**). In contrast, metaFlye and nanoMDBG could not recover any *P. copri* contigs of length *>* 700 kbp (less than 17% of the genome) (**Fig. 6B**).

**Fig. 6:**
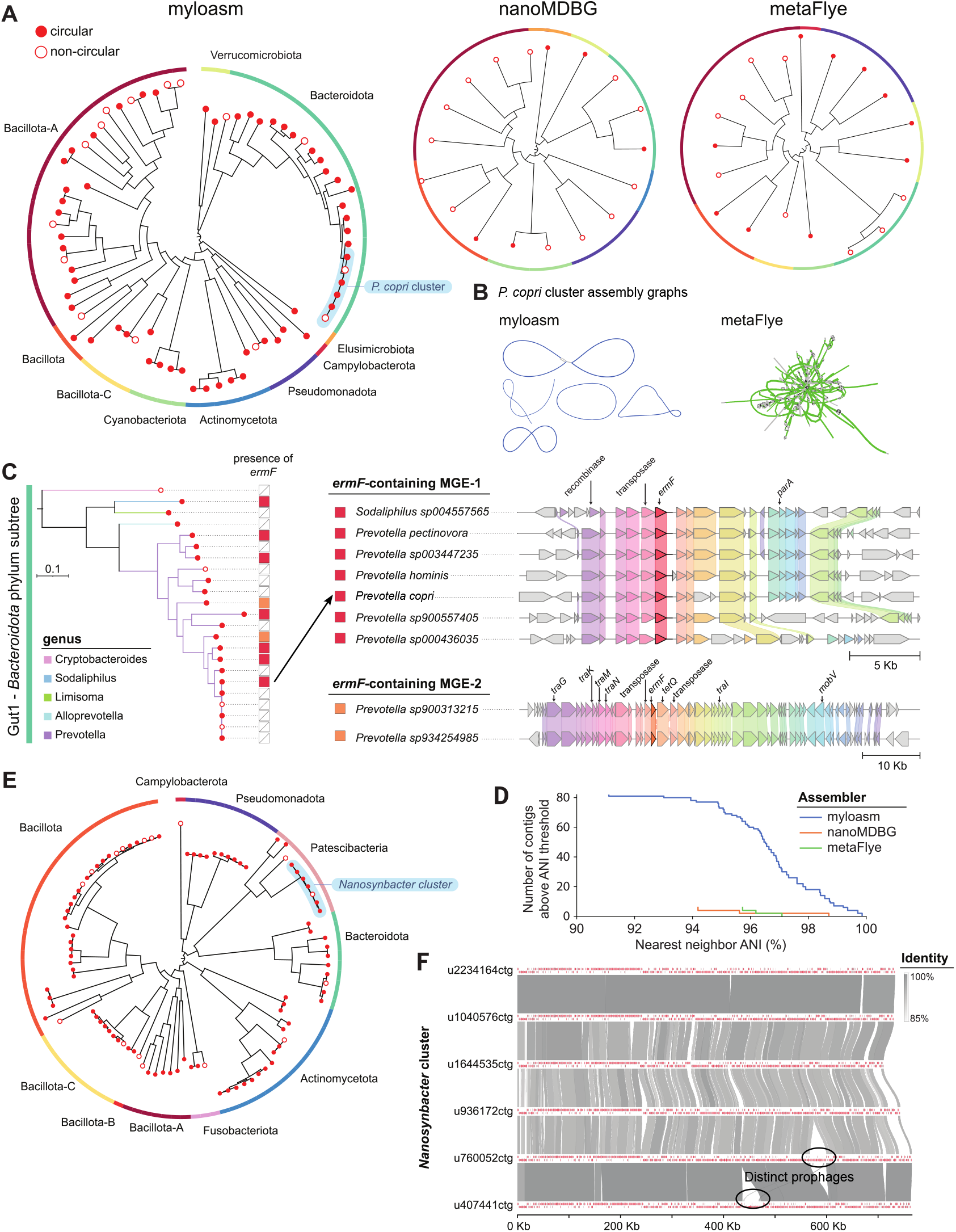
**A**. Phylogenetic of high-quality contigs (*>* 90% completeness and *<* 5% contamination) for the Gut 2 ONT sample. Phyla are labelled for myloasm’s tree, with six *P. copri* genomes outlined. **B**. Assembly graphs of *P. copri* for metaFlye and myloasm. *P. copri* contigs *>* 100 kbp are colored blue or green. **C**. Left: subtree for the Bacteroidota phylum. The right bar indicates the presence of an *ermF* gene, with color corresponding to the specific *ermF* sequence. Right: visualization of *ermF* gene contexts across contigs for the two distinct *ermF* genes. **D**. Number of contigs of length *>* 500 kbp with >90% ANI and *>* 70% AF to another contig within the assembly. **E**. Phylogenetic tree of high-quality contigs for Oral 1 ONT sample. **F**. Sequence alignments for six circular *Nanosynbacter* genomes recovered from the Oral 1 dataset.

High-contiguity assemblies enable investigations into mobile genetic elements and antibiotic resistance genes (ARGs) that short reads have difficulty assembling [63,31,64]. We highlight a particular example of *ermF*, an ARG that confers resistance to erythromycin through rRNA methylation [65]. *ermF* was prevalent within 9 of myloasm’s 23 Bacteroidota single-contig HQ genomes (**Fig. 6C**). Interestingly, only one of the six *P. copri* genomes had the *ermF* gene present, highlighting intraspecific patterns of ARG dissemination. The nine *ermF* genes could be separated into two distinct *ermF* sequences with 98% nucleotide identity to each other. Seven of the nine genomes shared an identical (100% nucleotide similarity) *ermF* sequence, confirming recent transfer across even Muribaculaceae and Bacteroidaceae families. Two of the nine genomes shared the other *ermF* sequence identically. These two distinct *ermF* genes could also delineated by their gene contexts; the second *ermF* sequence was clearly present in an integrative and conjugative element found in both *Prevotella* genomes. Thus, myloasm’s increased assembly resolution can resolve strain-specific patterns of genome dynamics and distinct evolutionary trajectories of antibiotic resistance genes.

Lastly, we investigated the Oral 1 ONT metagenome (**Fig. 6D-F**) to analyze a non-gut sample with high diversity. For all contigs of length ≥ 500 kbp, we analyzed how many contigs had *>* 90% ANI and >70% AF to another contig within the assembly (**Fig.6D**). Myloasm recovered 82 such contigs versus only 6 for nanoMDBG and metaFlye (**Supplementary Fig**. 5). An interesting example was a cluster of eight Saccharibacteria (also known as TM7) genomes of the genus *Nanosynbacter* with size ≈ 700 kbp (**Fig. 6E, F**). Six of the eight genomes were circular, but all had *>* 95% completeness (except one with 87.5%), *<* 0.6% contamination, and *>* 93.9% ANI to each other. We visualized the six circular *Nanosynbacter* genomes, finding perfectly conserved synteny even though several pairs of genomes had ANI *<* 95%, confirming previous findings on synteny conservation in TM7 [66]. We also found large unique regions within the *Nanosynbacter* genomes that had phage tail and capsid proteins, indicating the presence of strain-specific prophages. Thus, myloasm’s assemblies follow previous biological findings while highlighting the diversity across similar species and strains.

## 3 Discussion

We have created myloasm, a new metagenome long-read assembler that for modern long reads with greater than ≈ 97% accuracy, such as hac and sup base-called ONT R10.4 data and PacBio HiFi. Myloasm obtains substantially more MAGs and circular, high-quality contigs than previous ONT methods on synthetic and real data (**Fig. 2** and **Fig. 3**). Myloasm also often improves results for HiFi data compared to previous methods, and although the HiFi results are more comparable, it performs well for diverse metagenomes and sequencing technologies. Interestingly, myloasm demonstrates that ONT and HiFi reads are becoming more comparable, with myloasm even assembling more circular and complete genomes with ONT than HiFi (**Fig. 3B**) for a jointly-sequenced gut metagenome.

In general, our philosophy was to design myloasm as a metagenome assembler rather than adapt an isolate assembler for metagenomes. This led to our technical innovation of using SNPmers and SNPmer-associated overlap statistics. SNPmers are essentially a way of aggregating sequencing redundancy to find polymorphic markers, similar in spirit to error correction of reads. The ability to estimate true sequence identity through SNPmers gives a continuous measure of overlap similarity for overlaps and calculating coverage, which is key for metagenome assembly; the graph cleaning step makes substantial use of our coverage estimation process.

Myloasm substantially improves upon previous ONT assemblers when similar genomes are present (**Fig. 1** and **Fig. 6**), although this is not the only reason for myloasm’s improved assemblies (e.g., human gut with relatively low within-species diversity; see **Fig. 6A**). The term “strain-resolved” assembly is sometimes used to imply a method that can assemble multiple genomes of the same species [67,68,69]. There are fundamental issues to strain-resolved assembly, both theoretically and practically. One issue is that the notion of a clonal strain does not make sense in communities with high rates of recombination and gene flow. Theoretically, it is not even clear what a genome assembly means in this case, and we are not aware of existing mathematical theories that model genome assembly for continuous “cloud” of genotypes. A more practical issue is that even if we could assemble a collection of genomes that differ by a few substitutions, it may be preferable to output a single genome instead for interpretability. As sequencing technologies enable even higher resolution assemblies, these fundamental issues deserve more thought.

Finally, we note that errors are present in all assemblies [59]. Automated quality control software, such as CheckM2 and CheckV, are still necessary for contigs (**Fig. 4, 5**). We have found errors to arise due to a range of issues, from sequencing artifacts (e.g., chimeric reads; see **Methods**) to erroneous graph resolution for regions with high similarity. We provide additional scripts with the myloasm software for manually inspecting contigs via integrated plots of GC content, cumulative GC skew [70], read coverage, and read overlap lengths to assist users. Nevertheless, more work is needed to understand the algorithmic causes of assembly errors in complex metagenome assemblies [71]. We conclude by noting that long-read metagenome assembly is far from a solved problem, and we believe many opportunities still exist for optimization from both theoretical and practical points of view.

## 4 Methods

### 4.1 Capturing within-sample polymorphism with SNPmers

We use SNPmers in analogy to how heterozygous alleles are used to phase reads for diploid or polyploid genomes [72]. This is done by focusing on polymorphic sites between inter or intra-genomic repeats. Ignoring non-polymorphic bases increases the signal-to-noise ratio under sequencing error. However, since we do not have a reference genome, we use k-mers as a reference-free context [73,42] to call polymorphisms. Then, we compare these polymorphic k-mers to disambiguate strains.

#### Definition 1.

*Assume k, the k-mer length, is odd. A pair of k-mers is called a SNPmer pair if they have a different middle base but exact matches for the two flanks of length* (*k* − 1)*/*2. *For example, AACAA and AAGAA are a SNPmer pair for k* = 5. *We call any of the k-mers within a SNPmer pair a SNPmer. We only consider biallelic SNPmer pairs*.

We use *k* = 21 as the default parameter in myloasm. This choice is inspired by previous k-mer comparison methods that show *k* = 21 leads to a good balance between sensitivity and k-mer uniqueness in bacterial genomes [74].

### 4.2 Counting k-mers and filtering for SNPmers

To obtain a set of SNPmer pairs (and thus, SNPmers) for a sample, myloasm first counts all k-mers within the reads. We represent each k-mer in its canonical form: the lexicographically smaller of itself or its reverse complement. We keep track of whether the k-mer was in forward or reverse orientation relative to its canonical form. We ignore k-mers where the first (*k* − 1)*/*2 bases and the last (*k* − 1)*/*2 bases are reverse complements of each other. Such pairs are strand-ambiguous if the middle base is ignored. After obtaining all forward and reverse counts of each canonical k-mer, we retain only k-mers that have been seen once on both strands; we use a two bloom filters to filter out rare k-mers—one for each strand. This reduces erroneous SNPmers because sequencing errors often have strand bias [75].

To efficiently find SNPmer pairs, we lexicographically sort the canonical k-mers while masking the middle base. This groups all k-mers with the same flanking k-mers in the sorted list. If there are more than two k-mers with the same flanking (*k* − 1)*/*2-mers, we take the top two omst frequent k-mers as a potential SNPmer pair.

Finally, we retain these SNPmer pairs if they pass two additional tests. First, we test for strand bias by using a two-sided Fisher’s exact test on the 2×2 contingency table formed by the forward and reverse counts of the k-mer pair. We reject this pair if the odds ratio is *>* 1.5 and the p-value is *<* 0.005. Second, we use a binomial test based on the total count for each k-mer and a null error probability of 0.025: letting *A* be the total count for the higher frequency k-mer, and *B* the smaller frequency k-mer, we reject the SNPmer pair if *Pr*(*Binom*(*A*, 0.025) *> B*) *>* 0.05. If both tests pass, we retain the SNPmer pair.

### 4.3 Open syncmer and SNPmer read indexing

Myloasm indexes the reads with SNPmers from the previous step. However, SNPmers themselves are not reliable enough to map reads: genomes with minimal variation within the metagenome should have minimal SNPmers. Thus, we also index reads with *open syncmers* [44].

#### Definition 2.

*Given s < k, a k-mer is an open syncmer if the following holds: the smallest s-mer within the k-mer—subject to some fixed s-mer ordering—is the s-mer in position* (*k* − *s* + 1) *from the left (i*.*e*., *the middle s-mer). Define c* = (*k* − *s* + 1).

Under appropriate stochasticity assumptions, 1*/*(*k* − *s* + 1) = 1*/c* k-mers are sampled as open syncmers [44]. This leads to a speedup and memory improvement of *c* times. We order the s-mers by its numerical value after applying an invertible hash for 64-bit integers [47] and let *k* = 21 and *s* = 11 by default. We use open syncmers as opposd to the more common minimizer paradigm [76]. Open syncmers do not have the *context-dependency* property of minimizers [76,77]—minimizers lead to biased sequence identity estimates [78] that will impact our method later on. Furthermore, open syncmers have good empirical sensitivity [77,79] and theoretical guarantees for sequence alignment [80].

### 4.4 Read mapping and overlapping with double k-mer chaining

Define *anchor* to be a k-mer “match” (either SNPmer or open syncmer) between two sequences. These anchors are found in *different* ways: we let open syncmer anchors be *exact* k-mer matches. In contrast, we let SNPmer anchors be matches on *only the two flanking* (*k* − 1)*/*2*-mers*, thus ignoring the middle base. After finding all anchors, we perform two rounds of chaining: first on the open syncmers, then on the SNPmers.

#### Definition 3.

*We represent an anchor as a pair* (*x, y*) *of k-mer positions in the first and second sequence respectively. A co-linear chain (hereafter shortened to just chain) is a sequence of anchors* (*x*_1_, *y*_1_), (*x*_2_, *y*_2_), … *such that x*_*i*+1_ *> x*_*i*_ *and y*_*i*+1_ *> y*_*i*_. *Chaining is the computational problem of finding an optimal chain subject to some objective function*.

Chains capture the backbone of a sequence alignment and can be “extended” to form a full alignment. Note that to handle reverse complements, we can change the second condition to *y*_*i*_ *> y*_*i*+1_, but for exposition we will assume the forward case. For all stages except the last polishing stage, myloasm does not do any base level alignment, only chaining.

Algorithmically, we follow the chaining procedure in minimap2 [46] with modifications. Given a lexicographically sorted list of anchors, define the maximum chaining score for the *i*th anchor (*x*_*i*_, *y*_*i*_) as

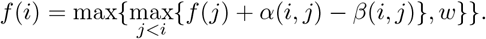

The goal of chaining is to to find the maximal *f* (*i*) over all anchors. By keeping track of the optimal predecessor *j* for each *i*, we can recover the chain that corresponds to the maximal *f* (*i*). *w* reflects the goodness of k-mer matching. *α*(*i, j*) = min{*x*_*i*_ − *x*_*j*_, *y*_*i*_ − *y*_*j*_, *w*} reflects the score of appending an additional anchor after penalizing overlapping anchors. *β*(*i, j*) = |(*x*_*i*_ − *x*_*j*_) − (*y*_*i*_ − *y*_*j*_)| is the gap cost. Additionally, we let *β* = −∞ if |*x*_*i*_ − *x*_*j*_| *> G* or |(*x*_*i*_ − *x*_*j*_) − (*y*_*i*_ − *y*_*j*_)| *> G*^*′*^ to break chaining for long gaps. We let *w* = *c* for open syncmer chaining (*c* = *k − s* + 1; *k* = 21 and *s* = 11 by default) and *w* = 50 for SNPmer chaining. *G*^*′*^ = 200 and *G* = 10000 by default.

Given *N* anchors, the optimal chain can be found in in *O*(*N* ^2^) time by computing *f* (*i*) with two for-loops and then backtracking. However, this quadratic time complexity is prohibitive. We therefore use minimap2’s heuristic of breaking the inner for-loop if the chaining score is not improved after *h* = 10 iterations.

### 4.5 Estimating true sequence alignment identity with double chains

After double chaining, myloasm checks the middle bases in the SNPmer chain. The number of mis-matched SNPmer anchors (i.e., mismatched middle bases) is due to both true differences and sequencing errors. Long-read sequencing errors are mostly indels [81]. Indels usually change the flanking (*k* − 1)*/*2-mers—SNPmer anchors would rarely occur under indels. On the other hand, true mutations for prokaryotes are usually substitutions [82]. Thus, SNPmer anchors should form for true mutations. Of course, substitution sequencing errors could lead to SNPmer anchors, but low-frequency substitutions errors should be filtered out by the SNPmer calling step. In addition, we enforce a Q23 base quality threshold (99.5% accuracy) on the middle base of a SNPmer match.

To summarize, we use the following intuition to estimate a true overlap sequence identity. (1) Sequencing errors will lead to *less anchors* (both SNPmer and open syncmer anchors), whereas (2) true mutations leads to SNPmers with *mismatched middle bases*. We use this intuition to estimate true sequence divergence through a probability model as follows.

#### Definition 4.

*Let X be a uniformly random string. Let Y be a version of X where each base is mutated to another base independently with probability θ. After mutation, mark each base as “erroneous” with probability ϵ*.

We examine the set of k-mer matches between *X* and *Y*. We exclude k-mers with an “erroneous” base under this idealized error model as follows. A k-mer on *X* is a mismatched SNPmer on *Y* with probability (1 − *θ*)^*k*−1^*θ* · (1 − *ϵ*)^*k*^: the middle base mutates (probability of *θ*), the other bases do not (probability of (1 − *θ*)^*k*−1^), and there are no sequencing errors (probability of (1 − *ϵ*)^*k*^). By linearity of expectation,

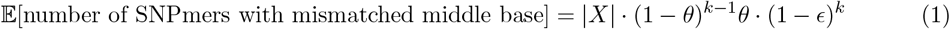

where |*X*| is the length of *X* (we assume that |*X*| ≈ |*X*| − *k* + 1 = the number of k-mers).

The left-hand side of Equation 1 can be estimated from SNPmer chaining. However, we can not estimate the true sequence identity *θ*: there is latent error *ϵ*. This is where we use the open syncmer chain. The probability of an open syncmer anchor forming is 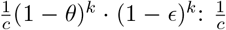 k-mers are open syncmers, and for the open syncmer to match, there must be no errors and no mutations. Note that this would not hold for minimizers [76] due to the minimizer context dependency issue [44,77,78]. Thus,

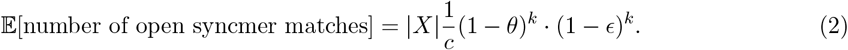

To estimate *θ*, we divide the two expressions.

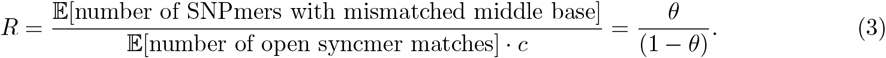

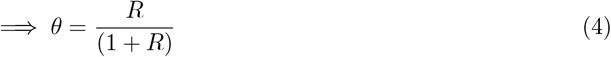

In summary, we can “divide out” the sequencing error by using the double chain. It turns out that the above idea leads to a *provably* consistent estimator for *θ* as the overlap length goes to ∞, under some mild assumptions.

#### Theorem 1.

*Let M*_*n*_ *be the number of mismatched SNPmers for a random string X and Y of length n under our random model. Let O*_*n*_ *be the analogous random variable for the number of matching open syncmers. Set*

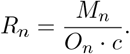

*Under an idealized model of X where every k-mer and s-mer is unique, as n* → ∞,

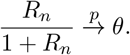

*That is, the left hand side converges to the right hand side in probability as the overlap gets large*.

We prove this theorem in the Supplementary Materials. We use techniques from Spouge et al. [83] showing that *M*_*n*_ and *O*_*n*_ have central limit theorems. This leads to our estimator of sequence identity, *θ*, as follows.

#### Definition 5.

*For a double chain between two sequences, let M be the number of mismatched SNPmers and O be the number of open syncmer anchors. Define*

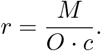

*Then, myloasm’s estimator of sequence identity*, 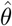, *is defined as*

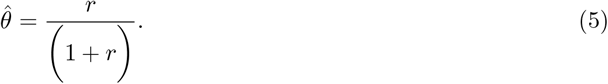

We show the effectiveness of the 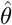 estimator in **Supplementary Fig. 6** relative to using a raw sequence alignment identity. Thus, 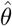 is an “error-corrected” divergence signal that allows us to disentangle distinguish error from true sequence divergence.

### 4.6 Removing contained reads

To build a string graph [35], we must remove reads that are contained (in another longer read). We will call a non-contained read an *outer read*. Removal of contained reads can be done by all-to-all read mapping. However, this *O*(*N* ^2^) computation for *N* reads is computationally intensive and often the main bottleneck in assembly workflows [84]. We instead devise a method that, for each queried read, quickly obtains a set of possible outer reads. As soon as the queried read is contained in *a single* read, we can move to the next query. This obviates the need for further alignments involving this query read.

Myloasm first builds a sparse hash table index with k-mers. For each read, myloasm filters out open syncmers if their hash value is *>* 2^64^ − 1)*/C* (*C* = 4 by default). This downsamples open syncmers by *C* = 4 times. Myloasm then inserts the filtered open syncmers as a key and appends its read ID to the value.

Next, for each query read, we query the hash table with all open syncmers. We track reads in the index that are “hit” by the query read’s open syncmers. We also track the query read’s leftmost and rightmost open syncmers for each read that is hit. After querying all open syncmers, we retain each hit read if the following hold: we require the rightmost and leftmost open syncmers of the query read— against a specific hit read—span 90% of the query read’s length. We then reverse sort these outer reads by the length of this span multiplied by the total number of open syncmers hit.

Finally, we do double chaining against each candidate outer read in the sorted list of hit reads. If (1) the number of mismatched SNPmers is 0 (i.e., 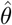 in Equation 5 is 0) and (2) the read open syncmer chain plus 30 · *c* spans 95% of the query read, then we remove the query read as a contained read.

### 4.7 Mapping to outer reads and calculating coverage

After removing contained reads, we map all reads against outer (i.e., non-contained) reads. The set of outer reads is usually much smaller than the set of all reads, so myloasm builds a new open syncmer index but only on the outer reads. Myloasm then performs all-to-outer double chaining and retains mappings with 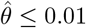.

Our goal is to calculate coverage for each read based on the read mappings. We differentiate two types of mappings: maximal mappings and local mappings.

#### Definition 6.

*Suppose the leftmost and rightmost positions for an open syncmer chain are* (*A*_1_, *B*_1_) *on read R*_1_ *and* (*A*_2_, *B*_2_) *on read R*_2_;. *let* |*R*| *be the length of a read R. A mapping is maximal if for some F (default = 300) the expression (A*_1_ *< F OR A*_2_ *< F) AND (B*_1_ + *F >* |*R*_1_| *OR B*_2_ + *F >* |*R*_2_|*) is true. Otherwise, it is a local mapping*.

Intuitively, a maximal mapping indicates that the mapping is as long as possible; it can not be extended much further without going past the end of a read. If the mapping from *R*_1_ to *R*_2_ is not maximal, there are regions without similar bases. Thus, the two reads are unlikely to originate from the same region of the same genome. Only maximal mappings should be used for calculating coverage.

We define the depth of coverage for each read as the *minimum* depth along each read, similar to Nurk et al. [85]. This is done to avoid overestimating coverage for repetitive regions. We define an identity-thresholded coverage as follows.

#### Definition 7.

*Let cov*_*R*_(*ω*) *be the minimum depth along R after removing maximal mappings with* 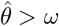.

We will calculate multiple coverages. We found that in high-diversity strain populations, there may be no perfect overlaps—that is, 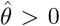 for every mapping. In this case, it is still useful to get a rough sense of coverage at lower resolutions.

### 4.8 Cleaning chimeric reads and large sequencing errors

Long reads can have large structural sequencing errors. For example, chimeric reads stitch together segments from two or more distinct genomes. Nanopore reads can also have large errors in low-complexity regions [86]. These errors can complicate the string graph. Long reads—especially ONT—can have highly heterogeneous lengths. Thus, long erroneous reads can eliminate contained reads and break contiguity [87,88]; see **Supplementary Fig. 7**. We found this to be the primary cause of errors in ONT assemblies.

We clean erroneous outer reads by using the all-against-outer local read mappings. If a region along the outer read has depth *d* ≤ 2 but one of the flanking regions 200 bp to the left or right has depth > 5 · *d*, then we break the read into two reads at this junction. For nanopore reads, if a region has depth *d* = 3 then we also break it if one flanking region have depth *>* 75.

After breaking chimeric outer reads, we repeat the containment and mapping procedure again, this time with the broken reads. This gives a second set of outer reads. Any remaining chimeric reads in the second round’s outer reads are broken one last time, and these outer reads are used for graph generation.

### 4.9 Read overlap graph generation with adaptive edge thresholds

Myloasm performs all-to-all read overlapping with double chaining for the final outer reads. We add an edge between two nodes if an overlap exists of length *>* 500 by default and 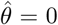; we discard reads of length *<* 1000. However, the requirement that 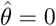 is sometimes too strict. It is possible that no reads come from the exact same strain or a sequencing error causes a SNPmer mismatch.

Thus, we use a coverage-informed threshold for 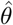 as follows. Recall the definition of *cov*_*R*_(*ω*) (Definition 7). If (1) *cov*_*R*_(0) *<* 5, (2) *cov*_*R*_(1*/*100) *>* 3 · *cov*_*R*_(0), and (3) *cov*_*R*_(1*/*100) *>* 5 then we define a new coverage threshold *ω*_*R*_. We use an iterative procedure to find *ω*_*R*_: first set *ω*_*R*_ = 0 and then iteratively calculate *cov*_*R*_(*ω*_*R*_ + 0.05*/*100). Stop when *cov*_*R*_(*ω*_*R*_ + 0.05*/*100) *>* 5 and 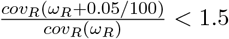. Finally, for an overlap between *R*_1_ and *R*_2_, we form an edge if 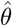 is less than 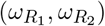.

We perform two more simplification procedures on this overlap graph.

#### Definition 8.

*Let the anchor sparsity of an overlap be the average bases between open syncmer anchors*.

We prune overlaps with anchor sparsity *>* 8 · *c*. Under a Bernoulli mutation model [89], this prunes overlaps of *<* 100 * (1*/*8)^1*/k*^ ≈ 90.6% sequence identity (k = 21). Given that modern nanopore reads have *>* 99% median accuracy [32], this did not cause many false negative overlaps. This heuristic was needed for the following reason: for very divergent overlaps, open syncmer chains could still form but no SNPmers were found, thus 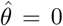. Mathematically, for higher sequence divergence *θ*, the *expected ratio* of mismatched SNPmers to syncmers may increase (Equation 3). But, the number of SNPmer mismatches still decreases (Equation 1) and is often 0 when *θ* is high.

We then prune overlaps with *>* 5 times higher anchor sparsity relative to the least sparse overlap.

We then rescue edges using two heuristics. We first rescue overlaps that are 1.5 times longer than the longest adjacent 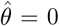 overlap and has 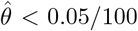. Second, we rescue overlaps with 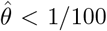 and connect tips to another node; we “rescue tips” if possible.

### 4.10 Bidirected string graph representation

Note that the initial read overlap graph is bidirected [35]. In a bidirected graph, an edge can be an in-edge or out-edge for either incident node. This represents the relative orientation of the overlapping reads. We define paths in a bidirected graph as follows.

#### Definition 9.

*For an edge e and an incident node x, let dir*(*e, x*) ∈ {*Out, In*} *be the incidence of e with respect to x. A path is a sequence of edges* (*e*_1_, *e*_2_, …, *e*_*n*_) *and nodes* (*x*_1_, *x*_2_, …, *x*_*n*+1_) *such that dir*(*e*_*i*−1_, *x*_*i*_) ≠ *dir*(*e*_*i*_, *x*_*i*_). *The length of a path is the length of the string it spells out*.

This definition naturally generalizes the notion of a path in a directed graph, where an edge *x* → *y* labelled *e* must have *dir*(*e, x*) = *Out* and *dir*(*e, y*) = *In*. However, this definition still works for a bidirected path: for example, (*e, e*^*′*^) for nodes (*x, y, z*) is a valid path if *dir*(*e, x*) = *In, dir*(*e, y*) = *In, dir*(*e*^*′*^, *y*) = *Out*, and *dir*(*e*^*′*^, *z*) = *Out*. Paths of overlapping reads—and thus, unitigs—spell out strings after appropriate reverse complementation.

### 4.11 Conservative unitig graph cleaning

We remove transitive edges in the cleaned overlap graph using the algorithm of Myers [35] and collapse non-branching paths into unitigs. This representation is the string graph, but we will refer to it as the unitig graph to make it clear that in subsequent cleaning steps, non-branching paths are always collapsed into unitigs.

We perform a set of conservative graph cleaning operations on the unitig graph. These operations requires a notion of *safety* for edge cutting. Intuitively, an edge is safe to cut if it does not break contiguity too badly. We illustrate this definition in **Supplementary Fig. 8**.

#### Definition 10.

*Let U*_1_ *and U*_2_ *be unitigs adjacent to an edge e. Let ℓ and L be safety two parameters. The edge e safe if the following are all true. (A) There exists a path* (*U*_*a*_, *x*, …) *with length > L where a* ∈ {1, 2} *and x* ≠ *U*_1_ *or U*_2_. *(B) For some path P* = (*U*_*a*_, *U*_*b*_, …, *x*_*i*−1_, *x*_*i*_, *x*_*i*+1_, …) *(where a, b* ∈ {1, 2}*) of length < ℓ, there exists a path* 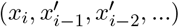 *that:*

1. *has length > L and*
2. *is edge-disjoint from P and*
3. *both x*_*i*−1_ *and* 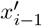 *are simultaneously in or out-nodes relative to x*_*i*_.

The first condition implies that cutting edge *e* does not create a “dead end”; the presence of an alternate path of length *> L* implies that cutting *e* does not break contiguity. The second condition implies that no dead ends will be created in a neighborhood of radius *ℓ* around *e*. Safety can be computed as follows. Path (A) in the definition can be found using a depth-first search (DFS). Path (B) can be found by doing a “forward” search of radius *ℓ* and then a “backward” DFS each time a new node is found for the forward search.

We cut safe edges using *ℓ* = 10kb and *L* = 50kb in two ways. These parameters will change later on. Firstly, we safely cut all edges that are *dominated* by another edge.

#### Definition 11.

*For a node x, an out-edge (resp. in-edge) e*_1_ *is dominated by another out-edge (resp. in-edge) e*_2_ *if has all* ***strictly*** *larger than e*_1_:

1. *overlap length*
2. *number of open syncmer anchors*
3. 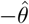
4. *number of matching SNPmer anchors*
5. −*number of mismatched SNPmer anchors*.

This heuristic removes edges that might have been erroneously rescued in the previous steps. Note that edges with 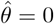 can not be dominated due to the strictness of the inequality.

We also cut safe edges with small relative overlap length as follows. Let *ol*(*e*) be the overlap length and *α <* 1. Let *e*_1_, *e*_2_, … be out or in-edges of some node. We cut *e*_*i*_ if for some *e*_*j*_ *ol*(*e*_*i*_) *< α* · *ol*(*e*_*j*_). We call this operation a drop cut.

Finally, we incorporate tip removal and bubble popping [47] with these graph cleaning procedures in an iterative fashion. First, we remove dominating edges. Let *α*, the drop cut ratio, increase from 0.5*/*3, 0.5*/*2, and then 0.5. In each iteration, we remove unitig tips that have length ≤ 20 kbp and ≤ 3 reads. Then, we pop bubbles of maximum length *<* 50 kb. Finally, we perform drop cuts with ratio *α*. After each step, we collapse new non-branching paths into unitigs. This corresponds to Lines 1-8 of Algorithm 1.

### 4.12 Path probability model

Recall that we can attach to each read a set of coverage values. In subsequent cleaning steps, we leverage coverage information. Intuitively true paths within the unitig graph should have consistent coverage because the sequencing coverage along a genome is roughly constant. We proceed by defining a probability model on paths in the graph. We subsequently use this model to weight edges. First, we define the coverage divergence between two unitigs.

#### Definition 12.

*Let U be a sequence of reads R*_1_, *R*_2_…. *For a threshold ω, the coverage of U is defined as an ordered set* 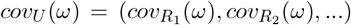; *see Definition 7. Let Q*_*x*_ *be the x-th percentile quantile function (e*.*g*., *x* = 50 *is the median). For U*_1_ *and U*_2_, *define*

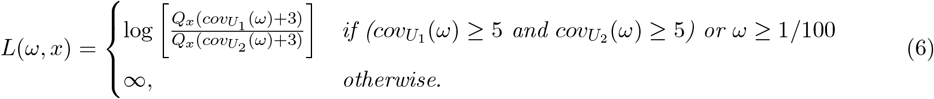

*Let Ω* = {1*/*100, 0.25*/*100, 0} *and define the coverage divergence between two sequences of reads be*

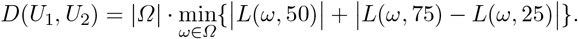

The first term of the coverage divergence is a log ratio of the median coverages. The second term captures differences in distributional shape. Myloasm stabilizes the *L*(*x, ω*) statistic by adding a pseudocount of 3 and only allowing a threshold *ω* if the coverage is high enough. This is crucial for strain populations where we often see *cov*_*R*_(0) ≈ 1 but *cov*_*R*_(1*/*100) ≫ 1. We found that adjacent unitigs can have highly variable coverages at specific values of *ω* due to e.g. inexact repeats, so the minimum smoothes out large variation. We now define the *energy* of a path.

#### Definition 13.

*Suppose we have a unitig path* (*U*_1_, …, *U*_*n*+1_) *with edges* (*e*_1_, …, *e*_*n*_). *Let Inc*(*U*_*i*_, *e*_*i*_) *be the in-edges (resp. out-edges) of U*_*i*_ *if e*_*i*_ *is an in-edge (resp. out-edge) of U*_*i*_. *Let* 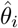 *be the* 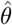 *for the ith edge and*

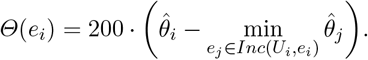

*Furthermore, let*

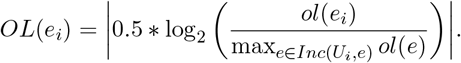

*Then the energy of a path is*

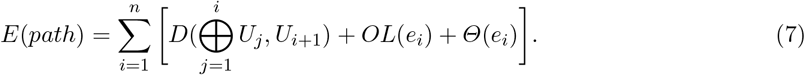

*where* 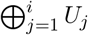 *is the concatenation of reads for all unitigs up to i*.

The definition of energy is arbitrary, but it encodes the intuition that we should penalize small overlaps, low sequence identity, and inconsistent converage (i.e., they should have high energy). We use the energy define a distribution over a class of paths as follows.

#### Definition 14.

*Assume some probability distribution over the vertices* Pr(*U*). *We choose*

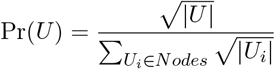

*where* |*U*| *is the number of reads in the starting unitig U in our implementation*.

*For a starting unitig node U, let* 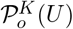 *be the set of out-paths from U that either have exactly K nodes or are not able to be extended further. Define* 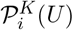 *similarly for in-paths. Then for* 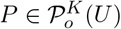, *and a “temperature” T >* 0 *let*

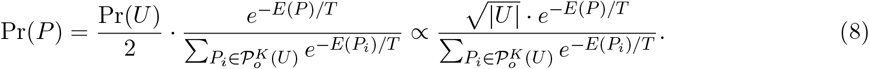

*If* 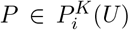, *define* Pr(*P*) *analogously. Note that a path traversed in the opposite direction is technically a different path (and thus may have different probability)*.

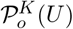 represent “maximal” paths from *U* of length at most *K* (see **Supplementary Fig. 8**). Intuitively, we think of generating these maximal paths of length ≤ *K* as follows. We first sample a node with probability Pr(*U*), then pick a direction with probability 1*/*2, and finally sample a path over all maximal length ≤ *K* paths from the node based on the Boltzmann distribution (or softmax function) of the path energies.

### 4.13 Path-infused edge weights

We use our path probability model to infuse edge weights with “global” information around a neighbourhood of the edge as follows.

#### Definition 15.

*Define the collection of paths* {*x* → *y*} *as the set of paths of the form* (…, *x, y*,). *Define* {*y* → *x*} *similarly. Note that* Pr({*x* → *y*}) ≠ Pr({*y* → *x*}); *the energy of a path depends on which direction it is traversed. Define the weight of an edge w*(*e*) *as*

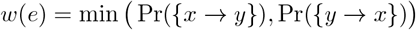

Intuitively, *w*(*e*) measures the likelihood of a local path flowing through the edge from both directions. We chose the minimum of the two directions because we often see nodes with skewed out-degree and in-degree in the graph (e.g., due to a conserved repeat at the end of one unitig).

#### Algorithm for computing edge weights

An important consideration is how to choose *K*, the number of nodes, in Definition 14. Depending on the genome, the string lengths spelled out by paths of length *K* can vary highly. Thus, we choose a *K*_*o*_(*U*) that depends on the starting node *U* and direction *o* or *i*. We set *K*_*o*_(*U*) to be the smallest number of nodes in an out-path path of length ≥ 1 Mbp (analogously for in-paths). If this is *>* 10, we set it to 10 instead. We discuss choosing the temperature *T* in the next section.

To actually compute the path probabilities in Definition 14, we would have to enumerate all paths. This is sometimes computationally prohibitive, so we use a beam search (i.e., top-N greedy breadth-first search) to prune out paths with low probability, as low-probability paths should not influence *w*(*e*) much. Precisely, we calculate the probability of each path by iterating over all nodes then running a beam search in both directions while keeping the N = 20 lowest energy paths at each iteration. Then, Pr({*x* → *y*}) is the sum of the probability of all paths passing from *x* to *y*.

### 4.14 Annealing-inspired graph cleaning

Once we have edge weights available, we cut safe edges (see Definition 10) with low edge weight relative to incident edges. That is, with *Inc*(*U, e*) defined as in Definition 13 and some *α*^*′*^ *<* 1, we cut edges with

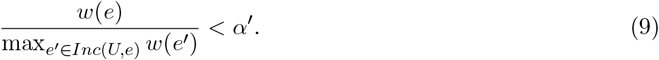

The crucial parameter for defining edge weights is the temperature *T*. *T* controls the strictness of the energy penalties in Definition 14. We circumvent a fixed choice of *T* by iterating through a range of *T* values in analogy to simulated annealing algorithms [41]. At high temperature, ratios of *w*(*e*) are closer to 1, so only very poor edges are cut. After cutting, re-unitigging yields larger unitigs with more reads. Subsequent calculations of energies in Definition 13 have larger “sample sizes” and thus higher reliability. Then, we repeat this procedure at lower *T*.

We summarize our entire graph cleaning procedure in Algorithm 1. Lines 9-20 correspond annealing-inspired cleaning step with additional tip removal and bubble popping. Notably, we also iterate on *α*^*′*^ and a parameter *m*. Recall that the definition of edge safety (Definition 10) depends on parameters *ℓ* and *L*—the larger *ℓ*, the less stringent the safety conditions. We make edge cuts more aggressive in Line 13 by multiplying the safety constant *ℓ* = 20000 by *m*. We also allow progressively larger bubbles of length *m* · 50 kb to be popped.

#### Algorithm 1 Graph cleaning pseudocode (G: initial unitig graph)

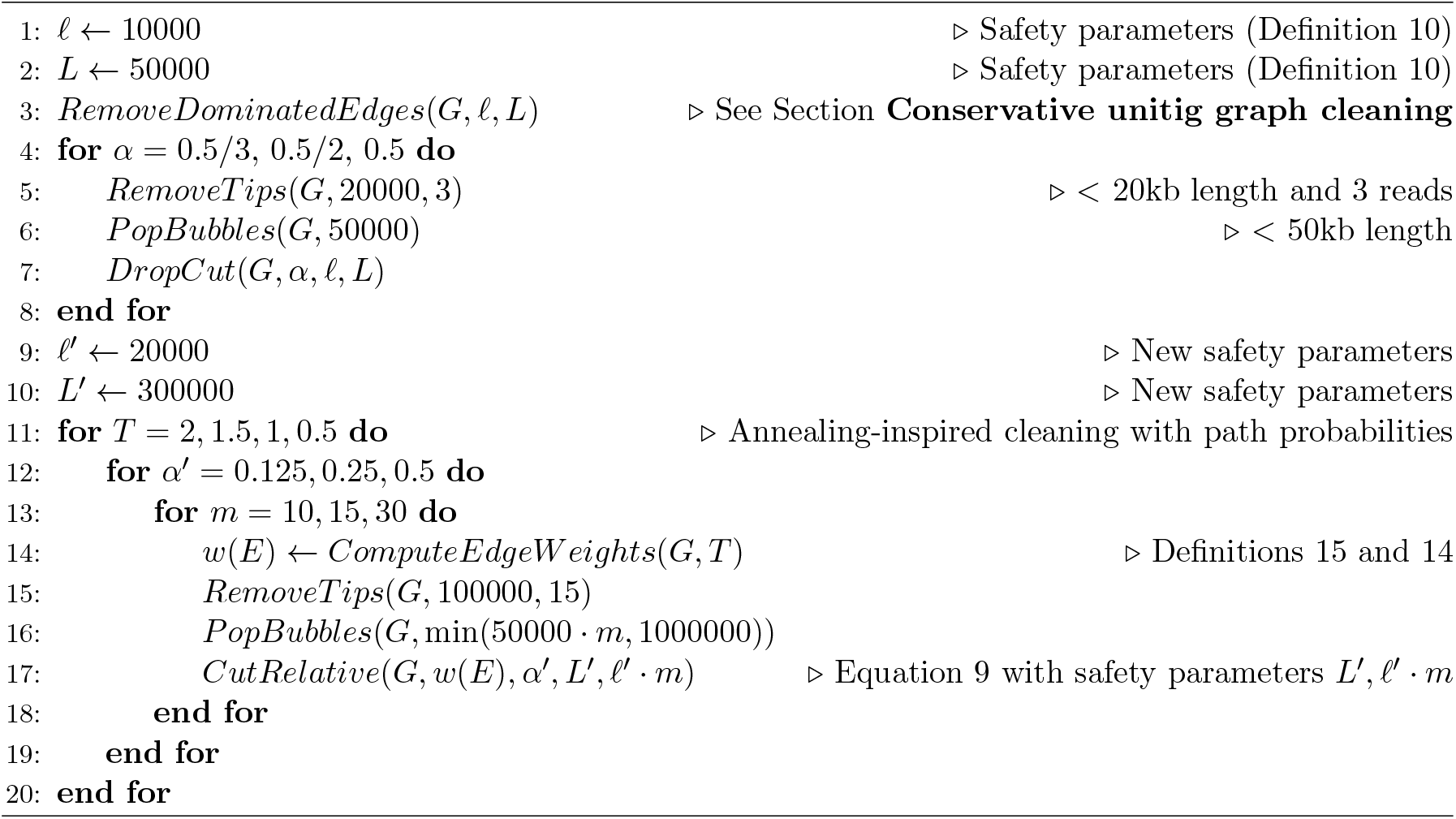

### 4.15 Circular contig extraction

We apply the progressive coverage filter of metaMDBG [20] for retrieving circular contigs as follows. We let cumulative coverage of a unitig be *cov*_*U*_ (0) + *cov*_*U*_ (0.25*/*100) + *cov*_*U*_ (1*/*100) (the same parameters as *Ω* in Definition 12). We remove contigs iteratively with cumulative coverage lower than *β* = 1, …, max(cumulative coverage) and re-unitig after each step. Afterwards, we initialize an empty set *C* = *∅* and iterate from *β*^*′*^ = max(cumulative coverage) 1. Suppose an isolated circular contig (i.e. in-degree = out-degree = 1) is found at an iteration. If the cumulative coverage of the circular contig is > 2 · *β*^*′*^ and none of its reads are in a contig in *C*, then we insert this contig into *C*. Finally, we remove all nodes in the original graph with a read in a contig of *C* and re-unitig. Then, we add all circular contigs in *C* back into the graph as isolated nodes.

Even after the above filter, very small circular plasmids were often not circularized properly. Thus, we use a heuristic extract small circular contigs as follows. If for a connected component in the final graph, every unitig has *<* 100 kb length and the entire component has *<* 100 reads, we mark it as a possible small circular genome. We recompute all pairwise overlaps and extract pairs of reads that form a putative circular unitig. Then, we sort pairs based on their estimated circular genome length. We iterate the through the sorted list until we find a genome that is 1.1 times longer than the previous genome length, taking the last pair of reads to be a circular unitig. The reason for this procedure is that we often found artifactual reads, e.g. reads from multimeric plasmids [49], that span the genome more than once. Sometimes pairs of these reads represent a repeated small genome, so we take the largest yet hopefully not repetitive pair of reads.

### 4.16 Polishing and dereplication

We first perform all-to-all pairwise mappings of unitigs and remove unitigs that are (1) *>* 98% contained in a larger unitig, (2) have 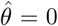, and (3) have *<* 8 · *c* anchor sparsity (that is, bases between anchors in the open syncmer chain on average; see Section 4.9 for justification).

We then align all reads to the remaining unitigs through double chaining. This time we extend the open syncmer chain to a full sequence alignment by performing global alignment between anchors using Block Aligner [90]. If a maximal mapping (see Definition 6) has *<* 8 · *c* anchor sparsity and 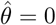, then we keep it. If a read contains no mapping that satisfies 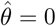, then we only retain a single best mapping scored by the number of open syncmer anchors + matching SNPmers - mismatched SNPmers.

After mapping, we polish using the SPOA (SIMD Partial Order Alignment) library [91,92] along 400bp windows to get the final contigs. We perform one last dereplication step by computing all-to-all average nucleotide identity (ANI) with skani [93] and the parameters `-c 15 -m 15`. We label contigs with *>* 99.9% ANI and *>* 99% fraction to a larger contig as duplicates and remove them. We label contigs with *>* 99% ANI and *>* 90% aligned fraction as alternates in a separate output.

### 4.17 Benchmarking setup

All assemblers were ran with default parameters except for technology-specific parameters. For metaFlye (v2.9.5-b1801), we used `--nano-hq` for nanopore and `--pacbio-hifi` for HiFi reads. For metaMDBG (v1.1), we used `--in-hifi` and `--in-ont` for HiFi and nanopore respectively. For myloasm (v0.1.0), we used `--hifi` for HiFi reads and default parameters otherwise. Hifiasm-meta (0.3-r074) was ran with default parameters. All assemblers were ran with 60 threads on a 64-core Intel(R) Xeon(R) Gold 6130 CPU machine with 384 GB of RAM except for the “Microflora” dataset [52] which required more memory. For this large complex soil dataset, we used a AMD EPYC 7301 machine with 1 TB of RAM. Timing and memory benchmarking was done through snakemake [94].

### 4.18 Mock synthetic R10.4 community benchmarking

We concatenated four separate datasets of ONT R10.4 reads with both hac and sup basecalled data available to form a mock metagenome. The three datasets were

1. Zymo Gut Microbiome Standard D6331 (dorado v0.8.2; v5.0.0 basecalling models)
2. Zymo Oral Microbiome Standard D6332 (dorado v0.8.2; v5.0.0 basecalling models)
3. Zymo HMW DNA Standard D6322 (dorado v0.7.3; v5.0.0 basecalling models)
4. 14 isolates from Hall et al. [32] (dorado v0.5.0; v4.3.0 basecalling models)

The zymo datasets were taken from https://github.com/Kirk3gaard/MicroBench. The fourteen isolates were downsampled to 10% of the original dataset unless the reads had *<* 1 Gbp, in which case we did not downsample. All datasets were concatenated and downsampled to 25% of the reads. This final dataset had 19 Gbp of data for 48 genomes after dereplicating reference genomes that had *>* 99.9% ANI according to skani. **Supplementary Table 3** shows the exact genome accessions and datasets used.

All assembly statistics were calculated by QUAST [95] as follows. We took all contigs of length *>* 100 kb for each assembler and matched it to the closest reference genome with skani. We ran QUAST on the set of matched contigs for each reference genome. We used this method as opposed to MetaQUAST [96] to have more control over strain-level assignments of contigs.

Qscore is defined as 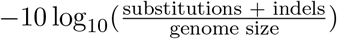 and was calculated from QUAST. We designated a contig as if the assembler designated it as circular and its length was *>* 1M bp. For plasmids, we did not use QUAST. Instead, we denoted a circular contig as a circular + perfect plasmid if skani (with parameter `--slow`) assigned it to a plasmid with *>* 99% ANI and both the contig and reference had >95% aligned fraction. If it satisfied the ANI and aligned fraction requirements but was not circular, we denoted it as a partial match. If the contig had *<* 90% aligned fraction to the plasmid but the plasmid had *>* 95% aligned fraction to the contig, then we designated it as a multimer and thus misassembled plasmid.

### 4.19 Real metagenome benchmarking procedure

We used six R10.4 nanopore datasets [51,50,52] and six PacBio HiFi datasets [20,53,56,54,55,57] and two samples sequenced with both HiFi and nanopore from Minich et al. Accessions are shown in Supplementary Table 1.

After assembly, we mapped reads back to the assemblies with minimap2 [46] and ran SemiBin2 [97] with default parameters for binning. We used CheckM2 [58] to calculate contamination and completeness of all resulting bins as well as contigs with length *>* 500 kb. We calculated zero-coverage windows within assemblies and positions with *>* 10 read clippings and no other alignments using a rust reimplementation of the `anvi-script-find-misassemblies` script from Anvi’o [98,59] available at https://github.com/bluenote-1577/rust-anvio-mis.

We used genomad [60] (v1.11.0) to find candidate viral and plasmid contigs. CheckV (v1.0.3) was used on all predicted viral contigs after excluding proviruses to assess quality. Conjugation and mobilization gene annotations were taken from genomad. To assess duplication, we obtained a k-mer frequency by counting all 21-mers in a contig and downsampling ≈ 1/10 k-mers with FracMinHash [99] and then finding the mean k-mer multiplicity after trimming the top 10% most and least-frequent k-mers. If this k-mer multiplicity was *>* 1.25, we marked the virus/plasmid contig as a duplicate.

We attempted to quantify the number of falsely circularized contigs, but ran into several issues. Firstly, simply quantifying the number of *<* 90% complete circular contigs is insufficient due to lineage-specific underestimation of genomic completeness in CheckM2. Furthermore, we found circularized contigs that were secondary chromosomes with low CheckM2 completeness. We also investigated *<* 500 kbp circular contigs with rRNA gene content but found several microeukaryotic organelle genomes in the environmental datasets (examples shown in **Supplementary Fig. 9**); furthermore, other types of small, circular extrachromosal elements can also have rRNA gene content [61].

### 4.20 Analysis of within-species diversity and horizontal gene transfer

All analysis of ANI and AF were done through skani with parameters `--medium --detailed --faster-small` `-m 400`. To construct the phylogenetic trees, we took all contigs of length *>* 500 kbp and completeness *>* 90% with contamination *<* 5%. We ran the `classify-wf` workflow of GTDB-TK [100] 2.3.2 with the GTDB-R214 taxonomy and then FastTree [101] with the GTDB-TK’s multiple sequence alignments of the `bac120` prokaryotic marker genes. Antibiotic resistance genes were annotated with the RGI pipeline using the CARD [102] database (May 2025 release). *De novo* gene annotation was performed on contigs individually using Bakta [103]. All circular contigs were updated to start at *dnaA* with the DNAapler [104] pipeline prior to Bakta annotation. Clinker [105] was used to visualize gene-level sequence relationships in genomic contexts surrounding predicted *ermF* locations in the Gut1-ONT dataset. The pgv-mummer workflow of pyGenomeViz (v1.6.1) was used to run mummer [106] iteratively between the six circular contigs of the nanosynbacter (CPR bacteria) cluster identified in the Oral1-ONT dataset.

## 5 Data availability

The mock nanopore community was generated from the MicroBench [107] suite by Rasmus Kirkegaard (https://github.com/Kirk3gaard/MicroBench and accession PRJEB85558) along with 14 isolates taken from Hall et. al [32]. Accessions for the real datasets are available in Supplementary Table 1.

## 6 Code availability

Myloasm is open source and available at https://github.com/bluenote-1577/myloasm. Documentation for myloasm is available at https://myloasm-docs.github.io/.

## 8 Acknowledgements

This work is supported by US National Institute of Health grant R01HG010040 to H.L. J.S. is supported by an NSERC Postdoctoral Fellowship (PDF) award. We thank members of the Li lab for helpful discussions.

## Supplementary Materials

### Proof of Theorem 1

#### Theorem 1.

Let *M*_*n*_ be the number of SNPmers with a mismatched middle base for a random string of length *X* of length *n*. Let *O*_*n*_ be the analogous random variable that counts the number of matching open syncmers. Set

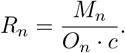

Under an idealized model of *X* where every k-mer and s-mer is unique, as *n* → ∞,

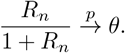

That is, the left hand side converges to the right hand side in probability.

*Proof*. Under an idealized random string model with k-mers and s-mers represented as uniformly random hashes, Spouge et al. [83] proved a central limit theorem (CLT) for “downsampled” (analogous to sequencing errors in our case) open syncmer matching:

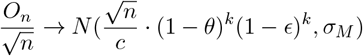

for some *σ*_*M*_.

Our strategy of proving Theorem 1 is as follows. Firstly, the above CLT implies that the law of large numbers as a corollary:

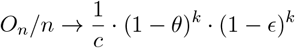

in probability. We will only need the law of large numbers, not the CLT.

We will show that

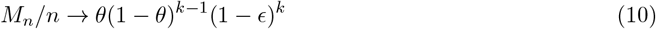

in probability by proving a CLT theorem for the SNPmers. Assuming this convergence holds,

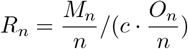

converges to

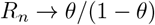

in probability by Slutsky’s theorem. Then the continuous mapping theorem with the function *x 1*→ *x/*(1+*x*) shows that *R*_*n*_*/*(1+*R*_*n*_) converges to *θ*, and we are done. Thus, our job is to prove Equation 10.

To prove Equation 10, we follow the same framework as Spouge et al. [83] and use a dependent central limit theorem of Hoeffding and Robbings [108] to prove that 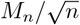 approaches a Gaussian.

#### Supplementary Lemma 1 (Hoeffding-Robbins CLT).

*Given a sequence of random variables Z*_1_, …, *Z*_*n*_, *this sequence is k-dependent if r* − *s > k implies that Z*_*i*+*r*_ *and Z*_*i*−*s*_ *are independent for any i. Assume that* 𝔼 [*Z*_*i*_] = 0 *and* 𝔼 [|*Z*_*i*_|^3^] *<* ∞. *Define*

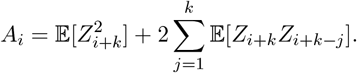

*Assume that* 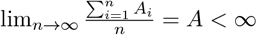 *exists. Then*, 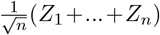 *has a limiting normal distribution with mean 0 and variance A*.

Along the random string *X*, our idealized SNPmers mismatches are *k*-dependent; this is because outside of this k-mer, the bases do not overlap and the letters are independent. Let *Z*_*i*_ = *z*_*i*_ − *p* where *p* = *θ*(1 − *θ*)^*k*−1^(1 − *ϵ*)^*k*^ and *z*_*i*_ is an indicator random variable that takes 1 if a SNPmer mismatch is found at position *i*. Clearly 𝔼 [*Z*_*i*_] = 0 by our previous discussion of 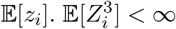 also holds because 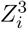 is an indicator random variable. We just need to show that 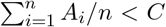 for some *C* and all *n*. The following inequality is sufficient:

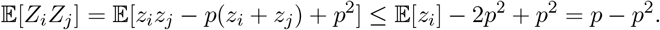

This holds because *z*_*i*_*z*_*j*_ ≤ *z*_*i*_ for all *i, j* (we are multiplying 0-1 indicators). Thus,

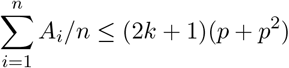

for all *n*, showing that 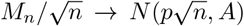 for some constant *A*. By CLT implying law of large numbers, this also proves Equation 10. This implication finishes the proof by the above argument.

**Supplementary Table 1:** Datasets for real ONT and HiFi datasets.

| **Dataset (ONT R10.4)** | **Accession** | **Dataset (HiFi)** | **Accession** |
| --- | --- | --- | --- |
| Oral 1 (Kiguchi et al.) | DRR582205 | Hot Spring (Kato et al.) | DRR290133 |
| Oral 2 (Kiguchi et al.) | DRR582179 | Anaerobic Digester (Benoit et al.) | ERR10905741 |
| Gut 1 (Minich et al.) | SRR29980972 | Chicken Gut (Zhang et al.) | SRR19683891 |
| Gut 2 (Minich et al.) | SRR29980959 | Human Gut 4 (Gehrig et al.) | SRR15489018 |
| Gut 3 (Minich et al.) | SRR29980980 | Seawater 1 (Priest et al.) | ERR4920901 |
| Microflora (Sereika et al.) | ERR11523665 | Seawater 2 (Sidhu et al.) | ERR9769281 |

**Supplementary Figure 1:**
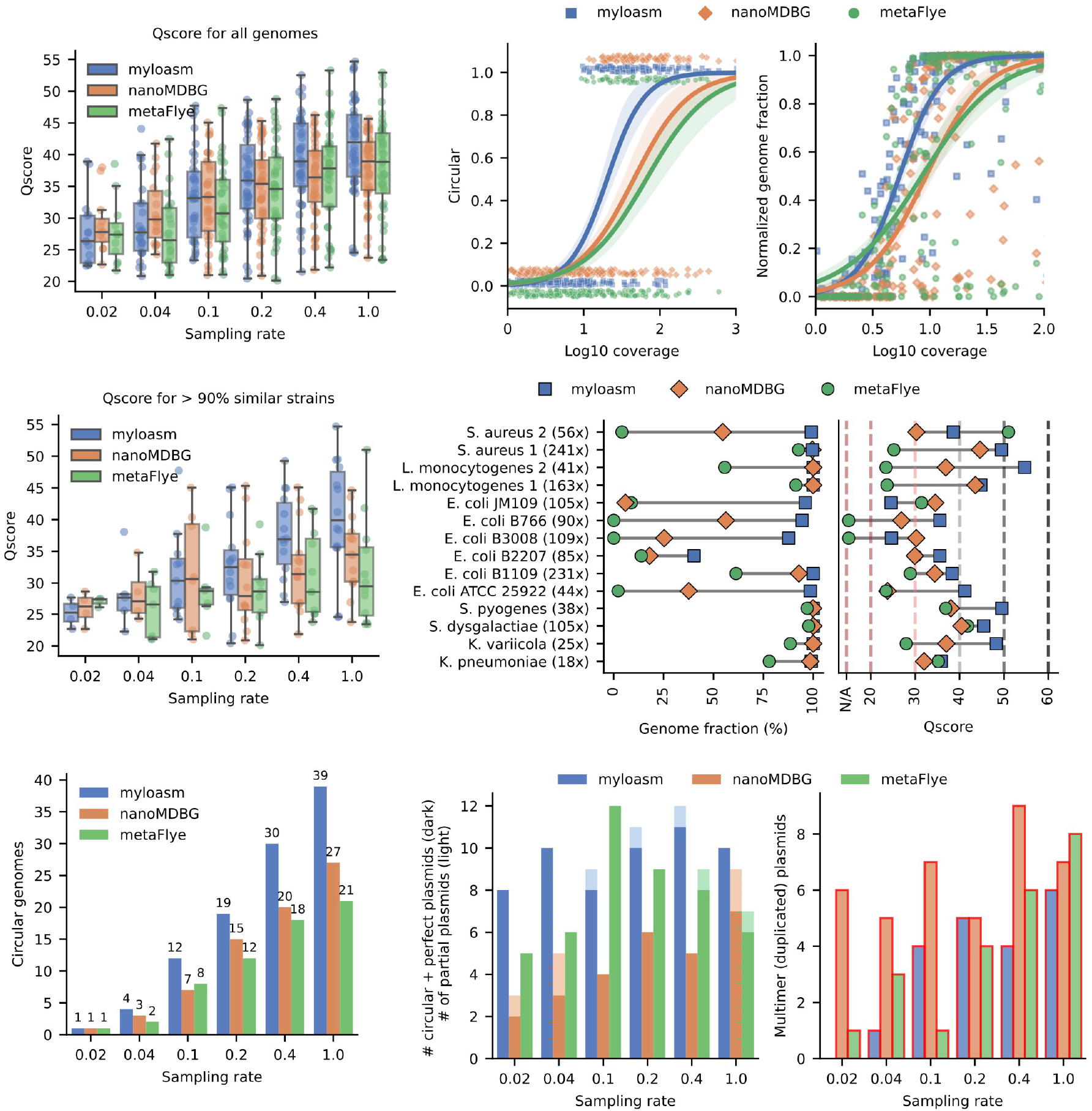
Results for the R10.4 ONT community but with hac (high accuracy) basecalling instead of sup (super high accuracy) basecalling. All subfigures are generated in the same way as in **Fig. 2**.

**Supplementary Figure 2:**
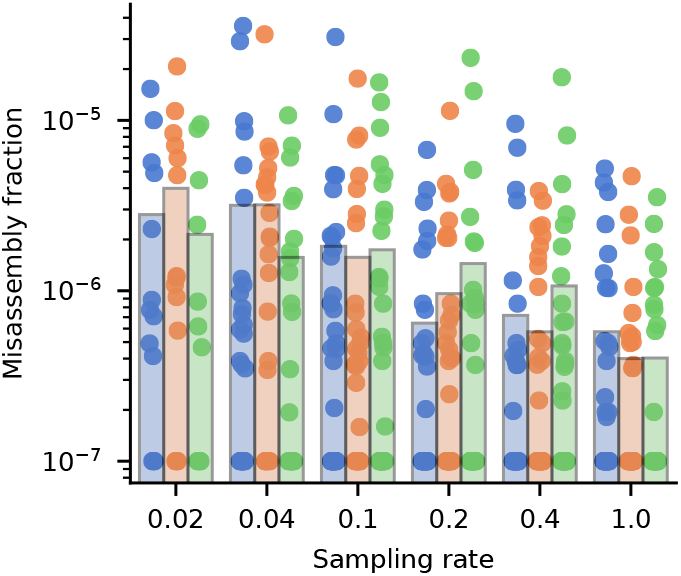
Misassembly fraction (# of misassemblies per base pair) over all genomes with Qscore *>* 20 for the mock metagenome (sup basecalling). Genomes without misassemblies were set to 10^−7^ misassembly fraction. The bar height represents the mean value.

**Supplementary Figure 3:**
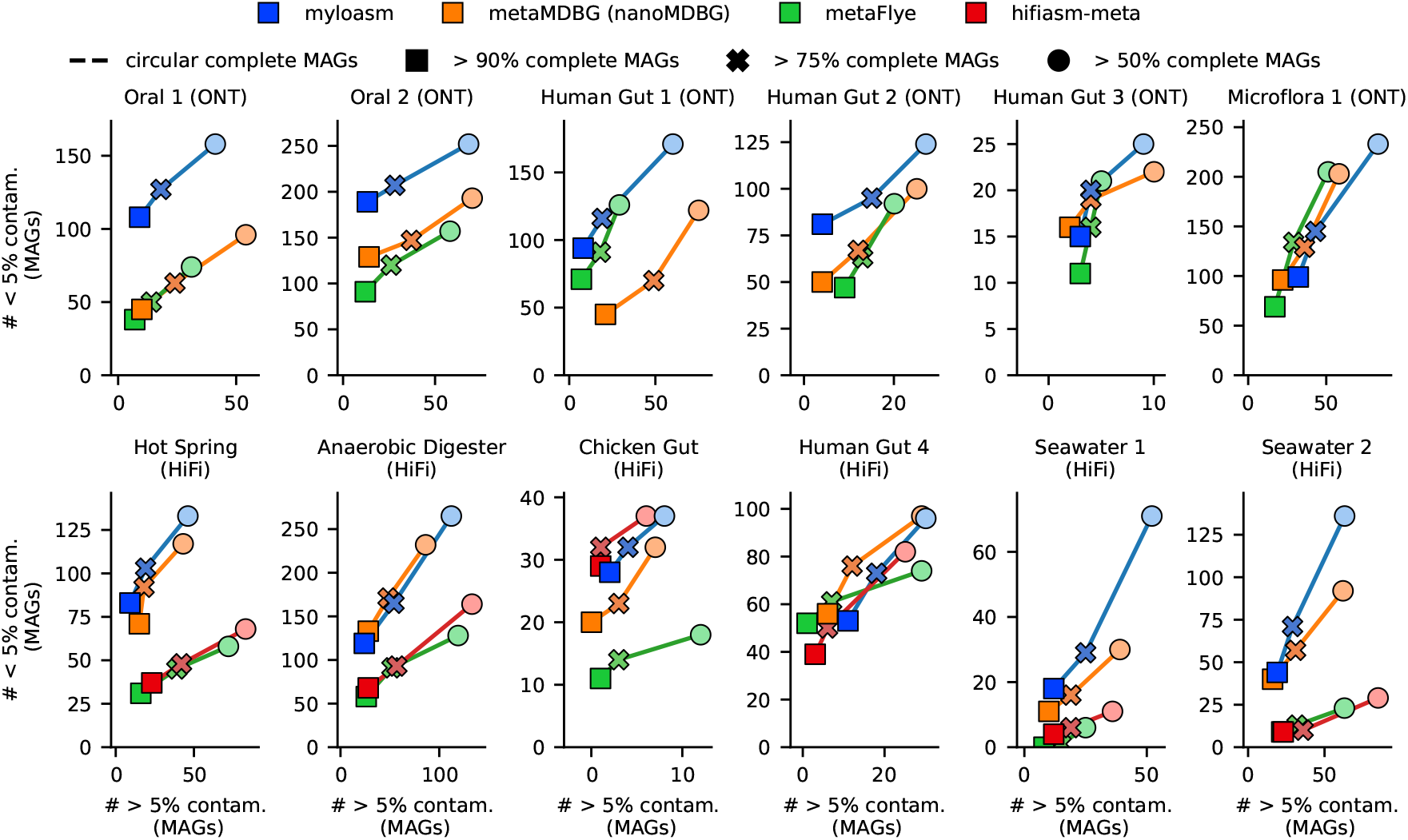
Analogous contamination and completeness results to **Fig. 4A** but for *MAGs*, not contigs.

**Supplementary Figure 4:**
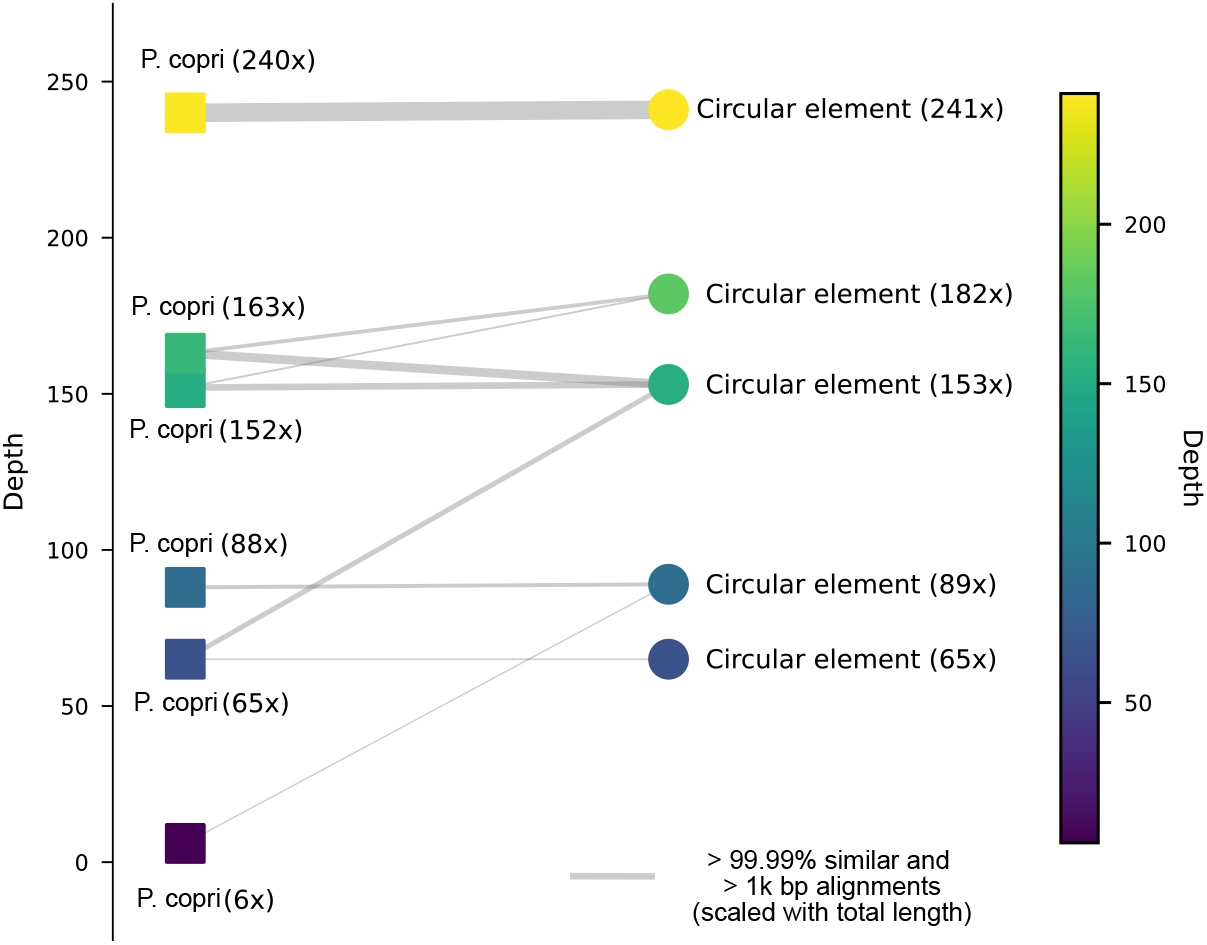
Linking *P. copri* genomes to circular genomic elements of length 100 − 150 kbp. Coverage was taken from myloasm’s estimated coverage (denoted as DP3 in the output). We confirmed the similarity of the 241x coverage circular contig against known extrachromosal elements of *P. copri* from Blanco-Miguez et al. [61]. We then found four more circular elements within the assembly with *>* 15% aligned fraction to the 241x element. Lines are drawn for matches of *>* 99.99% similarity as found by minimap2 between all circular elements and *P. copri* genomes, with thickness scaled linearly with total matching bases.

**Supplementary Figure 5:**
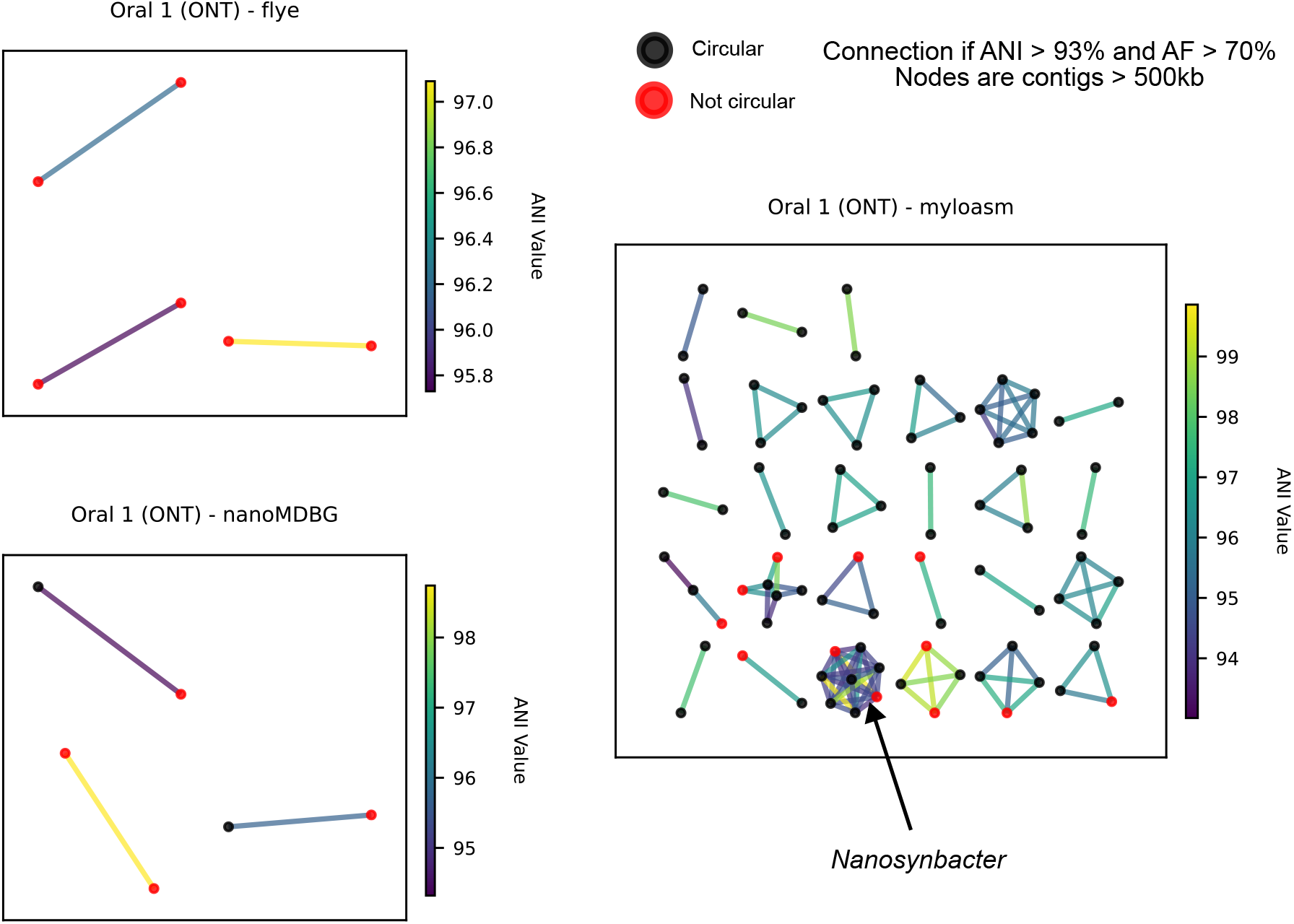
Contigs of length 500kbp in the Oral 1 dataset with at least one other contig with *>* 93% average nucleotide identity (ANI) and *>* 70% aligned fraction. Black dots are circularized genomes; red dots are non-circularized. Edges are drawn between genomes with *>* 93% ANI and *>* 70% aligned fraction and are colored by ANI.

**Supplementary Figure 6:**
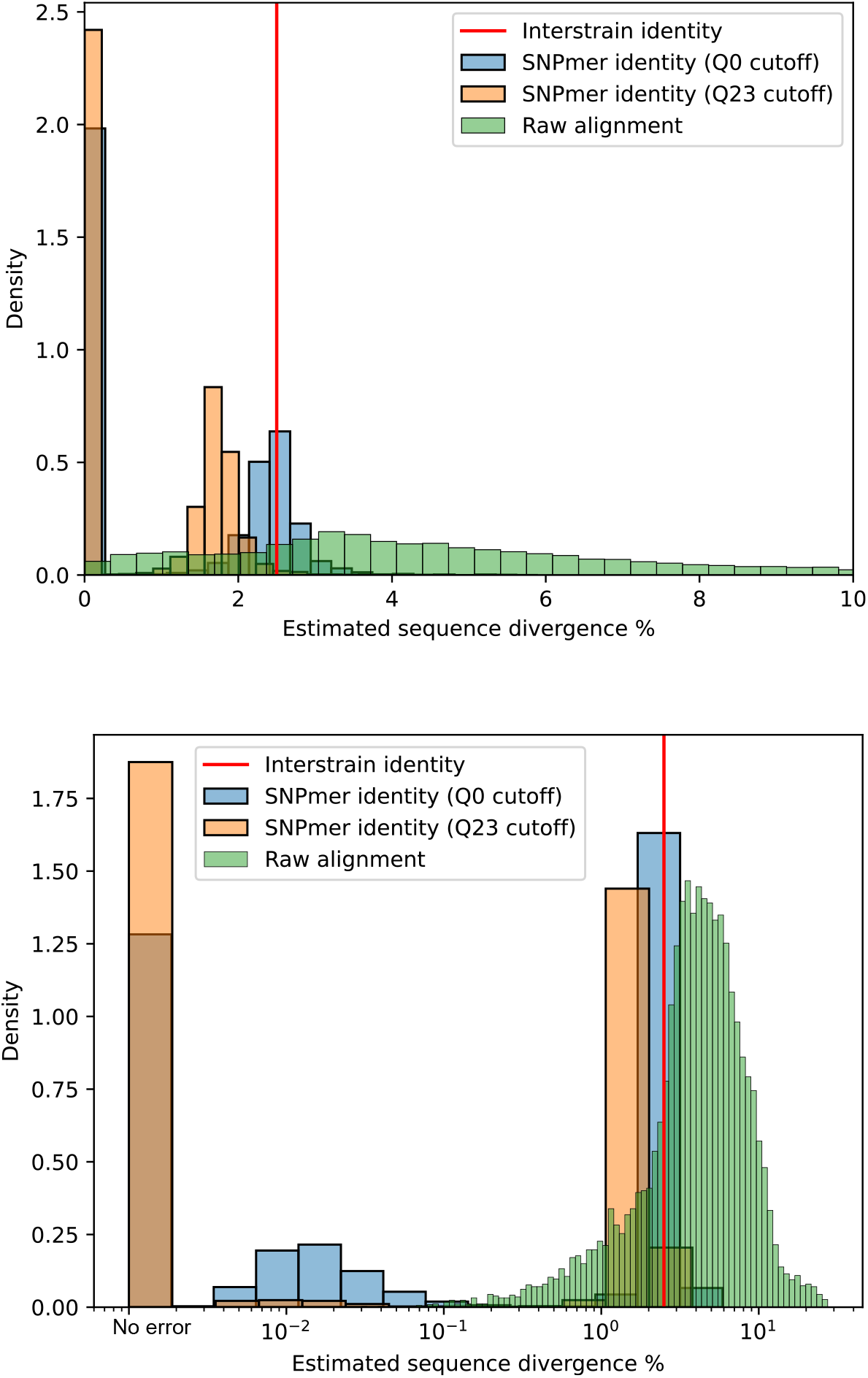
The SNPmer divergence 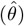 estimator (Equation 5) versus raw sequence identity for simulated reads from two *E. coli* genomes with 97.5% nucleotide similarity. Top: linear x-scale. Bottom: log x-scale. The second *E. coli* genome was generated by simulating substitutions with probability 0.025. Reads for both genomes were simulated with badread and default parameters except with mean accuracy 98%; the two read sets were then concatenated. Raw sequence identity was calculated by all-to-all read alignments of the strain-mixed reads with minimap2. We show two different values of 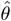: one after removing SNPmers with middle base quality *<* 23, and one with no base quality restrictions. Thresholding base qualities leads to a biased estimate of sequence divergence, but it also removes lots of low divergence overlaps (10^−2^% − 10^−1^%) that are from substitution sequencing errors (bottom).

**Supplementary Figure 7:**
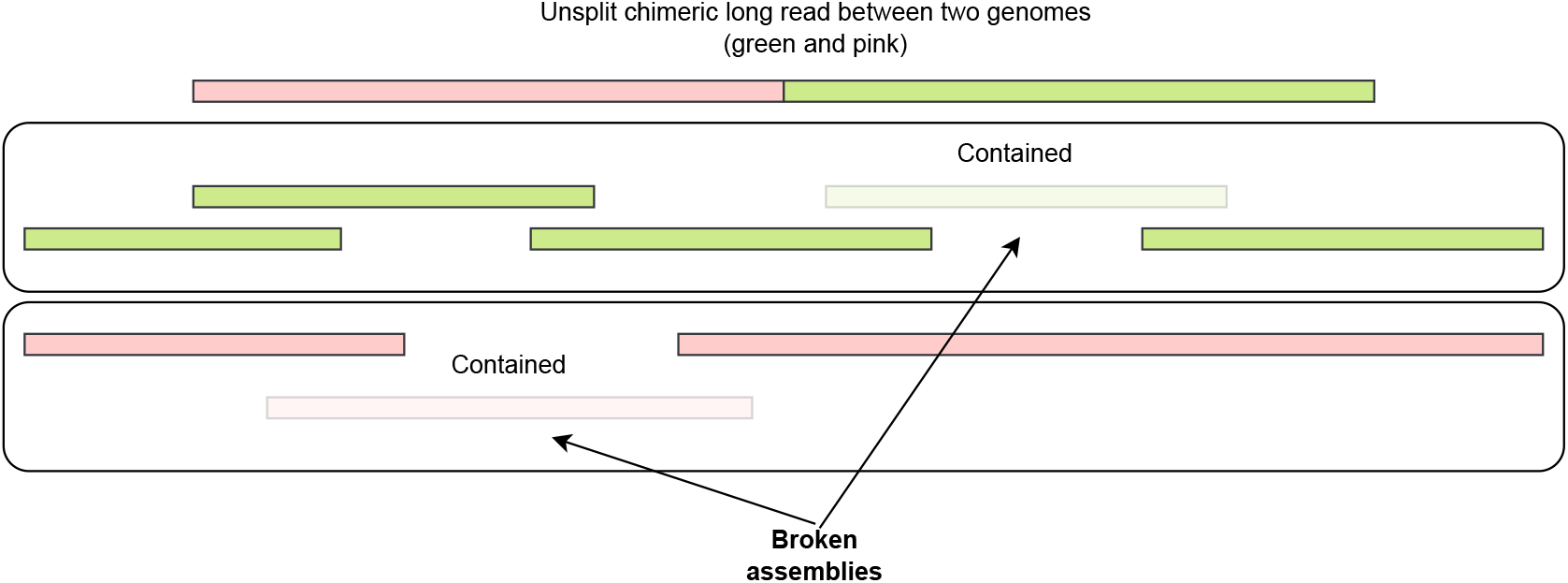
Illustration of how large structural errors within reads can cause misassemblies or fragmented assemblies. If a long chimeric read (top) is not split or removed, this read will remove smaller contained reads. The removal of the contained reads leads to broken assemblies, or the chimeric read may lead to a chimeric assembly.

**Supplementary Figure 8:**
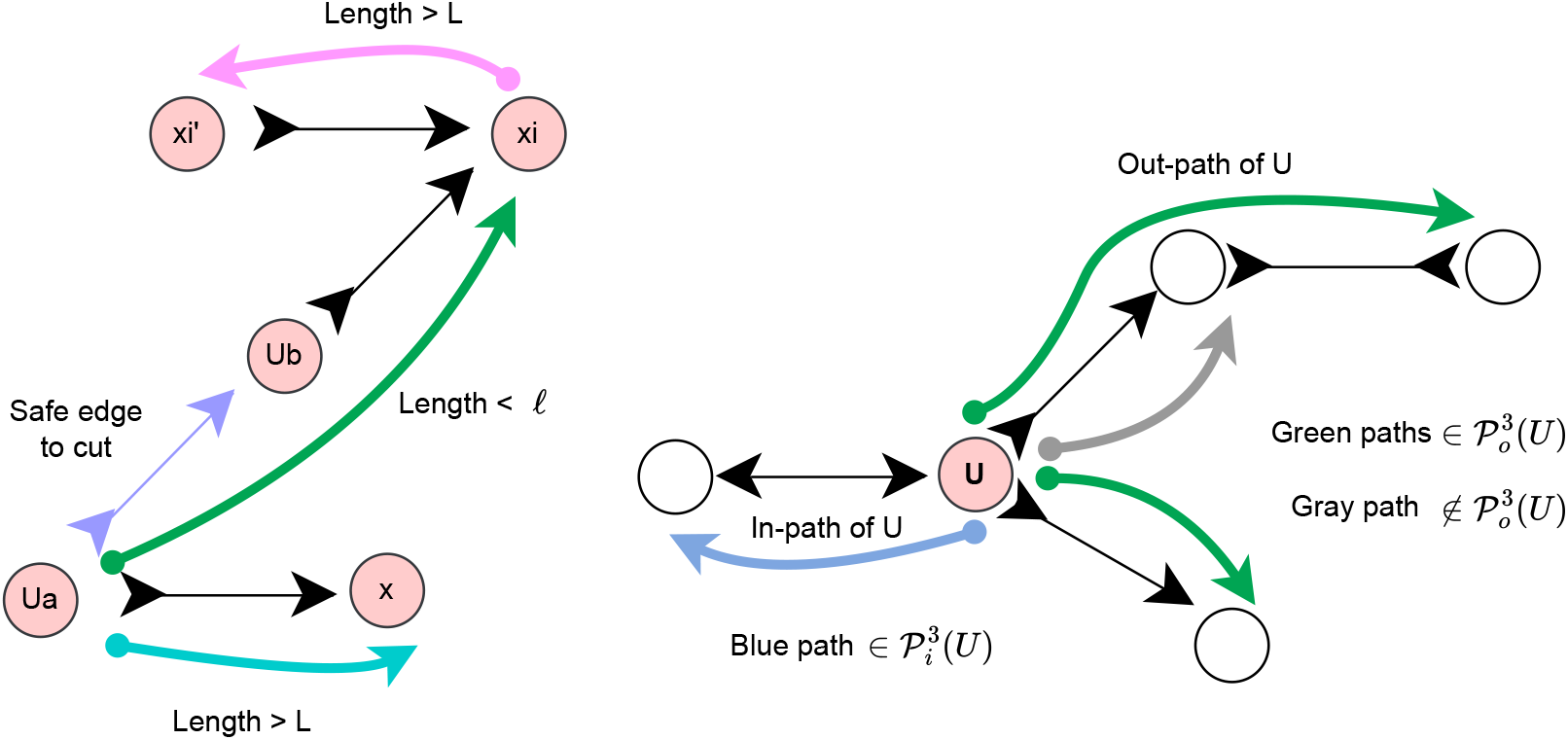
Illustration of key concepts in the bidirected graph representation of read overlaps. Edges in bidirected graphs have two independent directions—one direction for each node. **Left**: illustrating the definition of safety (Definition 10). The purple edge is safe to cut if there exists (1) a long path out of *U*_*a*_ (teal path) and (2) a long path (pink) that is “opposite” to a small path (green) out of *U*_*a*_. **Right:** illustrating Definition 14. In-paths and out-paths are shown. The gray path is not “maximal” for 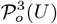 because it can be extended to a path of length 3. Thus, the gray path has no associated probability under our model.

**Supplementary Figure 9:**
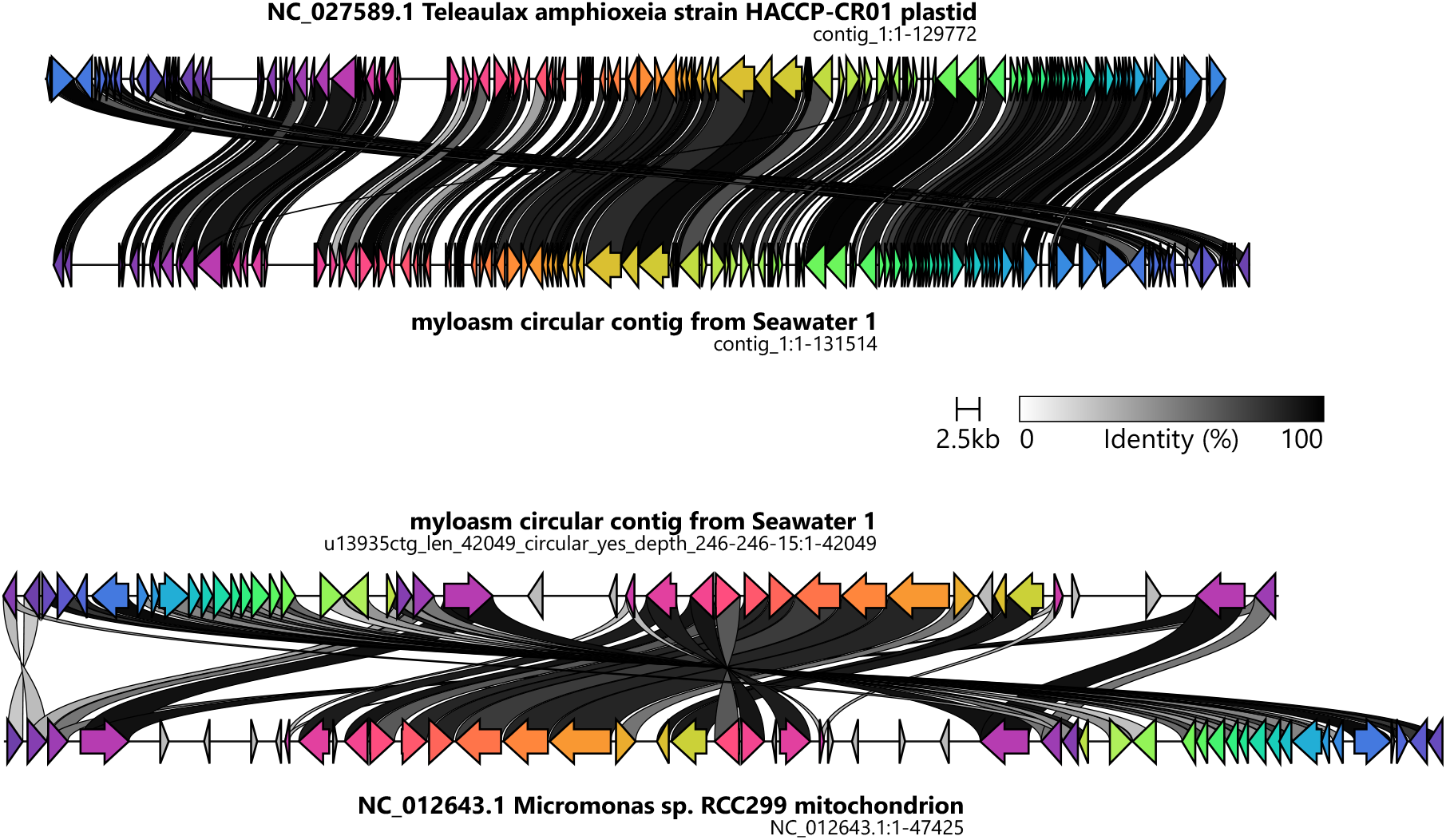
Circular and complete plastid and mitochondrial genomes recovered from the Seawater 1 dataset. The reference genomes NC_027589.1 and NC_012643.1 were the top hits after blasting the genomes to the NCBI blastn standard database. The reference genomes had 80 − 85% nucleotide similarity to myloasm’s contigs. Plots were generated by clinker [105].

## References

1. Quince, C., Walker, A. W., Simpson, J. T., Loman, N. J. & Segata, N. Shotgun metagenomics, from sampling to analysis. Nature Biotechnology 35, 833–844 (2017).

2. Hug, L. A. et al. A new view of the tree of life. Nature Microbiology 1, 1–6 (2016).

3. Parks, D. H. et al. Recovery of nearly 8,000 metagenome-assembled genomes substantially expands the tree of life. Nature Microbiology 2, 1533–1542 (2017).

4. Spang, A. et al. Complex archaea that bridge the gap between prokaryotes and eukaryotes. Nature 521, 173–179 (2015).

5. Zhao, S. et al. Adaptive Evolution within Gut Microbiomes of Healthy People. Cell Host & Microbe 25, 656–667.e8 (2019).

6. Pérez-Cobas, A. E., Gomez-Valero, L. & Buchrieser, C. Metagenomic approaches in microbial ecology: An update on whole-genome and marker gene sequencing analyses. Microbial Genomics 6, mgen000409 (2020).

7. Kiefl, E. et al. Structure-informed microbial population genetics elucidate selective pressures that shape protein evolution. Science Advances 9, eabq4632 (2023).

8. Wallen, Z. D. et al. Metagenomics of Parkinson’s disease implicates the gut microbiome in multiple disease mechanisms. Nature Communications 13, 6958 (2022).

9. Tisza, M. J. & Buck, C. B. A catalog of tens of thousands of viruses from human metagenomes reveals hidden associations with chronic diseases. Proceedings of the National Academy of Sciences 118, e2023202118 (2021).

10. Franzosa, E. A. et al. Gut microbiome structure and metabolic activity in inflammatory bowel disease. Nature Microbiology 4, 293–305 (2019).

11. Schmidt, T. S. B. et al. Drivers and determinants of strain dynamics following fecal microbiota transplantation. Nature Medicine 28, 1902–1912 (2022).

12. Bedarf, J. R. et al. Functional implications of microbial and viral gut metagenome changes in early stage L-DOPA-naïve Parkinson’s disease patients. Genome Medicine 9, 39 (2017).

13. Woodcroft, B. J. et al. Genome-centric view of carbon processing in thawing permafrost. Nature 560, 49–54 (2018).

14. Ustick, L. J. et al. Metagenomic analysis reveals global-scale patterns of ocean nutrient limitation. Science 372, 287–291 (2021).

15. Liang, J.-L. et al. Novel phosphate-solubilizing bacteria enhance soil phosphorus cycling following ecological restoration of land degraded by mining. The ISME Journal 14, 1600–1613 (2020).

16. Cavicchioli, R. et al. Scientists’ warning to humanity: Microorganisms and climate change. Nature Reviews Microbiology 17, 569–586 (2019).

17. Steen, A. D. et al. High proportions of bacteria and archaea across most biomes remain uncultured. The ISME Journal 13, 3126–3130 (2019).

18. Bertrand, D. et al. Hybrid metagenomic assembly enables high-resolution analysis of resistance determinants and mobile elements in human microbiomes. Nature Biotechnology 37, 937–944 (2019).

19. Kolmogorov, M. et al. metaFlye: Scalable long-read metagenome assembly using repeat graphs. Nature Methods 17, 1103–1110 (2020).

20. Benoit, G. et al. High-quality metagenome assembly from long accurate reads with metaMDBG. Nature Biotechnology (2024).

21. Feng, X., Cheng, H., Portik, D. & Li, H. Metagenome assembly of high-fidelity long reads with hifiasm-meta. Nature Methods 19, 671–674 (2022).

22. Agustinho, D. P. et al. Unveiling microbial diversity: Harnessing long-read sequencing technology. Nature methods 21, 954–966 (2024).

23. Feng, X. & Li, H. Evaluating and improving the representation of bacterial contents in long-read metagenome assemblies. Genome Biology 25, 92 (2024).

24. Crits-Christoph, A., Olm, M. R., Diamond, S., Bouma-Gregson, K. & Banfield, J. F. Soil bacterial populations are shaped by recombination and gene-specific selection across a grassland meadow. The ISME Journal 14, 1834–1846 (2020).

25. Liu, Z. & Good, B. H. Dynamics of bacterial recombination in the human gut microbiome. PLOS Biology 22, e3002472 (2024).

26. Chen-Liaw, A. et al. Gut microbiota strain richness is species specific and affects engraftment. Nature 637, 422–429 (2025).

27. Goyal, A., Bittleston, L. S., Leventhal, G. E., Lu, L. & Cordero, O. X. Interactions between strains govern the eco-evolutionary dynamics of microbial communities. eLife 11, e74987 (2022).

28. Brito, I. L. Examining horizontal gene transfer in microbial communities. Nature Reviews Microbiology 19, 442–453 (2021).

29. Nagarajan, N. & Pop, M. Parametric Complexity of Sequence Assembly: Theory and Applications to Next Generation Sequencing. Journal of Computational Biology 16, 897–908 (2009).

30. Bresler, G., Bresler, M. & Tse, D. Optimal assembly for high throughput shotgun sequencing. BMC Bioinformatics 14, S18 (2013).

31. Kerkvliet, J. J. et al. Metagenomic assembly is the main bottleneck in the identification of mobile genetic elements. PeerJ 12, e16695 (2024).

32. Hall, M. B. et al. Benchmarking reveals superiority of deep learning variant callers on bacterial nanopore sequence data. eLife 13, RP98300 (2024).

33. Sereika, M. et al. Oxford Nanopore R10.4 long-read sequencing enables the generation of near-finished bacterial genomes from pure cultures and metagenomes without short-read or reference polishing. Nature Methods 19, 823– 826 (2022).

34. Cheng, H. et al. Efficient near telomere-to-telomere assembly of Nanopore Simplex reads (2025).

35. Myers, E. W. The fragment assembly string graph. Bioinformatics (Oxford, England) 21 Suppl 2, ii79–85 (2005).

36. Compeau, P. E. C., Pevzner, P. A. & Tesler, G. Why are de Bruijn graphs useful for genome assembly? Nature biotechnology 29, 987–991 (2011).

37. Ekim, B., Berger, B. & Chikhi, R. Minimizer-space de Bruijn graphs: Whole-genome assembly of long reads in minutes on a personal computer. Cell Systems 12, 958–968.e6 (2021).

38. Benoit, G. et al. High-quality metagenome assembly from nanopore reads with nanoMDBG. bioRxiv 2025.04.22.649928 (2025).

39. Cheng, H., Concepcion, G. T., Feng, X., Zhang, H. & Li, H. Haplotype-resolved de novo assembly using phased assembly graphs with hifiasm. Nature Methods 18, 170–175 (2021).

40. Nurk, S. et al. HiCanu: Accurate assembly of segmental duplications, satellites, and allelic variants from high-fidelity long reads. Genome Research 30, 1291–1305 (2020).

41. Kirkpatrick, S., Gelatt, C. D. & Vecchi, M. P. Optimization by Simulated Annealing. Science 220, 671–680 (1983).

42. Derelle, R. et al. Seamless, rapid and accurate analyses of outbreak genomic data using Split K-mer Analysis (SKA) (2024).

43. Gardner, S. N. & Hall, B. G. When Whole-Genome Alignments Just Won’t Work: kSNP v2 Software for Alignment-Free SNP Discovery and Phylogenetics of Hundreds of Microbial Genomes. PLOS ONE 8, e81760 (2013).

44. Edgar, R. Syncmers are more sensitive than minimizers for selecting conserved k-mers in biological sequences. PeerJ 9, e10805 (2021).

45. Myers, G. & Miller, W. Chaining multiple-alignment fragments in sub-quadratic time. In Proceedings of the Sixth Annual ACM-SIAM Symposium on Discrete Algorithms, SODA ‘95, 38–47 (Society for Industrial and Applied Mathematics, USA, 1995).

46. Li, H. Minimap2: Pairwise alignment for nucleotide sequences. Bioinformatics 34, 3094–3100 (2018).

47. Li, H. Minimap and miniasm: Fast mapping and de novo assembly for noisy long sequences. Bioinformatics 32, 2103–2110 (2016).

48. Bouras, G. et al. Hybracter: Enabling scalable, automated, complete and accurate bacterial genome assemblies. Microbial Genomics 10, 001244 (2024).

49. Vaisbourd, E., Bren, A., Alon, U. & Glass, D. S. Preventing Multimer Formation in Commonly Used Synthetic Biology Plasmids. ACS Synthetic Biology 14, 1309–1315 (2025).

50. Minich, J. J. et al. Culture-independent meta-pangenomics enabled by long-read metagenomics reveals novel associations with pediatric undernutrition. SSRN Electronic Journal (Preprint) (2024).

51. Kiguchi, Y. et al. Giant extrachromosomal element “Inocle” potentially expands the adaptive capacity of the human oral microbiome. Nature Communications 16, 7397 (2025).

52. Sereika, M. et al. Recovery of highly contiguous genomes from complex terrestrial habitats reveals over 15,000 novel prokaryotic species and expands characterization of soil and sediment microbial communities (2024).

53. Gehrig, J. L. et al. Finding the right fit: Evaluation of short-read and long-read sequencing approaches to maximize the utility of clinical microbiome data. Microbial Genomics 8, 000794 (2022).

54. Sidhu, C. et al. Dissolved storage glycans shaped the community composition of abundant bacterioplankton clades during a North Sea spring phytoplankton bloom. Microbiome 11, 77 (2023).

55. Priest, T., Orellana, L. H., Huettel, B., Fuchs, B. M. & Amann, R. Microbial metagenome-assembled genomes of the Fram Strait from short and long read sequencing platforms. PeerJ 9, e11721 (2021).

56. Kato, S., Masuda, S., Shibata, A., Shirasu, K. & Ohkuma, M. Insights into ecological roles of uncultivated bacteria in Katase hot spring sediment from long-read metagenomics. Frontiers in Microbiology 13 (2022).

57. Zhang, Y. et al. Improved microbial genomes and gene catalog of the chicken gut from metagenomic sequencing of high-fidelity long reads. GigaScience 11, giac116 (2022).

58. Chklovski, A., Parks, D. H., Woodcroft, B. J. & Tyson, G. W. CheckM2: A rapid, scalable and accurate tool for assessing microbial genome quality using machine learning. Nature Methods 20, 1203–1212 (2023).

59. Trigodet, F., Sachdeva, R., Banfield, J. F. & Eren, A. M. Assemblies of long-read metagenomes suffer from diverse errors. bioRxiv 2025.04.22.649783 (2025).

60. Camargo, A. P. et al. Identification of mobile genetic elements with geNomad. Nature Biotechnology 42, 1303–1312 (2024).

61. Blanco-Míguez, A. et al. Extension of the Segatella copri complex to 13 species with distinct large extrachromosomal elements and associations with host conditions. Cell Host & Microbe 31, 1804–1819.e9 (2023).

62. Chang, H.-W. et al. Prevotella copri and microbiota members mediate the beneficial effects of a therapeutic food for malnutrition. Nature Microbiology 9, 922–937 (2024).

63. Maguire, F. et al. Metagenome-assembled genome binning methods with short reads disproportionately fail for plasmids and genomic Islands. Microbial Genomics 6, mgen000436 (2020).

64. Abramova, A., Karkman, A. & Bengtsson-Palme, J. Metagenomic assemblies tend to break around antibiotic resistance genes. BMC Genomics 25, 959 (2024).

65. Xing, L. et al. ErmF and ereD Are Responsible for Erythromycin Resistance in Riemerella anatipestifer. PLoS ONE 10, e0131078 (2015).

66. He, X. et al. Cultivation of a human-associated TM7 phylotype reveals a reduced genome and epibiotic parasitic lifestyle. Proceedings of the National Academy of Sciences of the United States of America 112, 244–249 (2015).

67. Kazantseva, E., Donmez, A., Frolova, M., Pop, M. & Kolmogorov, M. Strainy: Phasing and assembly of strain haplotypes from long-read metagenome sequencing. Nature Methods 1–10 (2024).

68. Shaw, J., Gounot, J.-S., Chen, H., Nagarajan, N. & Yu, Y. W. Floria: Fast and accurate strain haplotyping in metagenomes. Bioinformatics 40, i30–i38 (2024).

69. Jochheim, A. et al. Strain-resolved de-novo metagenomic assembly of viral genomes and microbial 16S rRNAs. Microbiome 12, 187 (2024).

70. Grigoriev, A. Analyzing genomes with cumulative skew diagrams. Nucleic Acids Research 26, 2286–2290 (1998).

71. Schmidt, S., Toivonen, S., Medvedev, P. & Tomescu, A. I. Applying the Safe-And-Complete Framework to Practical Genome Assembly. LIPIcs: Leibniz international proceedings in informatics 312, 8 (2024).

72. Lancia, G., Bafna, V., Istrail, S., Lippert, R. & Schwartz, R. SNPs Problems, Complexity, and Algorithms. In auf der Heide, F.M. (ed.) Algorithms — ESA 2001, Lecture Notes in Computer Science, 182–193 (Springer, Berlin, Heidelberg, 2001).

73. Chaung, K. et al. SPLASH: A statistical, reference-free genomic algorithm unifies biological discovery. Cell 186, 5440–5456.e26 (2023).

74. Ondov, B. D. et al. Mash: Fast genome and metagenome distance estimation using MinHash. Genome Biology 17, 132 (2016).

75. Liu, X. et al. Nanopore strand-specific mismatch enables de novo detection of bacterial DNA modifications. Genome Research 34, 2025–2038 (2024).

76. Roberts, M., Hayes, W., Hunt, B. R., Mount, S. M. & Yorke, J. A. Reducing storage requirements for biological sequence comparison. Bioinformatics 20, 3363–3369 (2004).

77. Shaw, J. & Yu, Y. W. Theory of local k-mer selection with applications to long-read alignment. Bioinformatics 38, 4659–4669 (2022).

78. Belbasi, M., Blanca, A., Harris, R. S., Koslicki, D. & Medvedev, P. The minimizer Jaccard estimator is biased and inconsistent. Bioinformatics (Oxford, England) 38, i169–i176 (2022).

79. Frith, M. C., Shaw, J. & Spouge, J. L. How to optimally sample a sequence for rapid analysis. Bioinformatics 39, btad057 (2023).

80. Shaw, J. & Yu, Y. W. Proving sequence aligners can guarantee accuracy in almost O(m log n) time through an average-case analysis of the seed-chain-extend heuristic. Genome Research gr.277637.122 (2023).

81. Delahaye, C. & Nicolas, J. Sequencing DNA with nanopores: Troubles and biases. PLOS ONE 16, e0257521 (2021).

82. Chen, J.-Q. et al. Variation in the Ratio of Nucleotide Substitution and Indel Rates across Genomes in Mammals and Bacteria. Molecular Biology and Evolution 26, 1523–1531 (2009).

83. Spouge, J. L., Das, P., Chen, Y. & Frith, M. The Statistics of Parametrized Syncmers in a Simple Mutation Process Without Spurious Matches. Journal of Computational Biology 31, 1195–1210 (2024).

84. Stanojević, D., Lin, D., de Sessions, P. F. & Šikić, M. Telomere-to-telomere phased genome assembly using errorcorrected Simplex nanopore reads. bioRxiv 2024.05.18.594796 (2024).

85. Nurk, S. et al. The complete sequence of a human genome. Science 376, 44–53 (2022).

86. Tan, K.-T., Slevin, M. K., Meyerson, M. & Li, H. Identifying and correcting repeat-calling errors in nanopore sequencing of telomeres. Genome Biology 23, 180 (2022).

87. Jain, C. Coverage-preserving sparsification of overlap graphs for long-read assembly. Bioinformatics 39, btad124 (2023).

88. Li, H. & Durbin, R. Genome assembly in the telomere-to-telomere era. Nature Reviews Genetics 25, 658–670 (2024).

89. Blanca, A., Harris, R. S., Koslicki, D. & Medvedev, P. The Statistics of k-mers from a Sequence Undergoing a Simple Mutation Process Without Spurious Matches. Journal of Computational Biology: A Journal of Computational Molecular Cell Biology 29, 155–168 (2022).

90. Liu, D. & Steinegger, M. Block Aligner: An adaptive SIMD-accelerated aligner for sequences and position-specific scoring matrices. Bioinformatics 39, btad487 (2023).

91. Vaser, R., Sović, I., Nagarajan, N. & Šikić, M. Fast and accurate de novo genome assembly from long uncorrected reads. Genome Research 27, 737–746 (2017).

92. Lee, C., Grasso, C. & Sharlow, M. F. Multiple sequence alignment using partial order graphs. Bioinformatics 18, 452–464 (2002).

93. Shaw, J. & Yu, Y. W. Fast and robust metagenomic sequence comparison through sparse chaining with skani. Nature Methods 1–5 (2023).

94. Köster, J. & Rahmann, S. Snakemake—a scalable bioinformatics workflow engine. Bioinformatics 28, 2520–2522 (2012).

95. Gurevich, A., Saveliev, V., Vyahhi, N. & Tesler, G. QUAST: Quality assessment tool for genome assemblies. Bioinformatics 29, 1072–1075 (2013).

96. Mikheenko, A., Saveliev, V. & Gurevich, A. MetaQUAST: Evaluation of metagenome assemblies. Bioinformatics 32, 1088–1090 (2016).

97. Pan, S.Zhao, X.-M. & Coelho, L. P. SemiBin2: Self-supervised contrastive learning leads to better MAGs for short-and long-read sequencing. Bioinformatics 39, i21–i29 (2023).

98. Eren, A. M. et al. Anvi’o: An advanced analysis and visualization platform for ‘omics data. PeerJ 3, e1319 (2015).

99. Rahman Hera, M., Pierce-Ward, N. T. & Koslicki, D. Deriving confidence intervals for mutation rates across a wide range of evolutionary distances using FracMinHash. Genome Research 33, 1061–1068 (2023).

100. Chaumeil, P.-A., Mussig, A. J., Hugenholtz, P. & Parks, D. H. GTDB-Tk: A toolkit to classify genomes with the Genome Taxonomy Database. Bioinformatics 36, 1925–1927 (2020).

101. Price, M. N., Dehal, P. S. & Arkin, A. P. FastTree 2 – Approximately Maximum-Likelihood Trees for Large Alignments. PLOS ONE 5, e9490 (2010).

102. Alcock, B. P. et al. CARD 2023: Expanded curation, support for machine learning, and resistome prediction at the Comprehensive Antibiotic Resistance Database. Nucleic Acids Research 51, D690–D699 (2023).

103. Schwengers, O. et al. Bakta: Rapid and standardized annotation of bacterial genomes via alignment-free sequence identification. Microbial Genomics 7, 000685 (2021).

104. Bouras, G., Grigson, S. R., Papudeshi, B., Mallawaarachchi, V. & Roach, M. J. Dnaapler: A tool to reorient circular microbial genomes. Journal of Open Source Software 9, 5968 (2024).

105. Gilchrist, C. L. M. & Chooi, Y.-H. Clinker & clustermap.js: Automatic generation of gene cluster comparison figures. Bioinformatics 37, 2473–2475 (2021).

106. Marçais, G. et al. MUMmer4: A fast and versatile genome alignment system. PLOS Computational Biology 14, e1005944 (2018).

107. Kirkegaard, R. MicroBench (https://github.com/Kirk3gaard/MicroBench) (2025).

108. Hoeffding, W. & Robbins, H. The Central Limit Theorem for Dependent Random Variables. In Fisher, N.I. & Sen, P.K. (eds.) The Collected Works of Wassily Hoeffding, 205–213 (Springer New York, New York, NY, 1994).

